# Visualizing nuclear pore complex plasticity with pan-Expansion Microscopy

**DOI:** 10.1101/2024.09.18.613744

**Authors:** Kimberly J. Morgan, Emma Carley, Alyssa N. Coyne, Jeffrey D. Rothstein, C. Patrick Lusk, Megan C. King

## Abstract

The exploration of cell-type and environmentally-responsive nuclear pore complex (NPC) plasticity requires new, accessible tools. Using pan-Expansion Microscopy (pan-ExM), NPCs were identified by machine learning-facilitated segmentation with resolved cytoplasmic rings (CR), inner rings (IR) and nuclear rings (NR). They exhibited a large range of diameters with a bias for dilated NPCs at the basal nuclear surface in clusters suggestive of local islands of nuclear envelope (NE) tension. Whereas hyperosmotic shock constricted NPCs analogously to those found in annulate lamellae (AL), depletion of LINC complexes specifically eliminated the modest nuclear surface diameter biases. Therefore, LINC complexes may contribute locally to nuclear envelope tension to toggle NPC diameter between dilated, but not constricted, states. Lastly, POM121 shifts from the NR to the IR specifically in induced pluripotent stem cell derived neurons (iPSNs) from a patient with *C9orf72* amyotrophic lateral sclerosis (ALS). Thus, pan-ExM is a powerful tool to visualize NPC plasticity in physiological and pathological contexts at single NPC resolution.

**Summary:** Morgan et al. demonstrate that pan-Expansion Microscopy is an exceptional tool for probing the molecular composition and structural plasticity of individual NPCs: they reveal LINC-complex dependent NPC diameter biases across the nuclear surface and changing nucleoporin positions in an ALS model.

## Introduction

The nuclear envelope (NE) is a double membrane barrier that segregates the nuclear genome from the cytoplasm. Bidirectional molecular communication across the NE is mediated by nuclear pore complexes (NPCs), protein gateways that facilitate the selective transport of cargo-bound nuclear transport receptors (NTRs) while also imposing a size-restrictive diffusion barrier. It is well understood how the majority of the key building blocks (nucleoporins or nups) are arranged to build the iconic 8-fold radial architecture of the NPC with recent cryo-EM and cryo-ET structures approaching atomic resolution (Akey et al., 2023; Akey et al., 2022; Bley et al., 2022; Fontana et al., 2022; Huang et al., 2022a; Huang et al., 2022b; Kosinski et al., 2016; Mosalaganti et al., 2022; Petrovic et al., 2022; Schuller et al., 2021; Singh et al., 2024; von Appen et al., 2015; Zhu et al., 2022). These studies have provided a blueprint for the NPC in multiple organisms, glimpses of its evolutionary history and key insights into its assembly and function. They have also revealed that the NPC structure is not monolithic. Indeed, there are numerous hints that NPCs are surprisingly plastic in composition and structure, but the causes and consequences of this likely plasticity are just beginning to come to light (Fernandez-Martinez and Rout, 2021). A key challenge for the field is to be able to directly visualize this plasticity on a single NPC level.

The NPC scaffold is built from three concentric ring assemblies, the cytoplasmic ring (CR), inner ring (IR) and nuclear ring (NR) that delimit the central transport channel. Recent work supports that the NPC scaffold constricts in response to energy depletion or hyperosmotic shock (Zimmerli et al., 2021) suggesting that lateral strain on NPCs imposed by NE tension modulates NPC architecture and function (Akey et al., 2022; Mosalaganti et al., 2022; Schuller et al., 2021). It remains unknown, however, whether changing the diameter of NPCs impacts their selective permeability (although there is emerging evidence that supports this concept)(Elosegui-Artola et al., 2017; Feng et al., 2024; Klughammer et al., 2024; Kozai et al., 2023). Further, it is unclear whether NPC diameter is controlled by passive mechanisms or whether it can be actively modulated, perhaps even in response to the local NE environment. Considering the latter, it is plausible that mechanotransduction mechanisms that translate extracellular mechanical cues to the NE through <u>Li</u>nker of <u>N</u>ucleoskeleton and <u>C</u>ytoskeleton (LINC) complexes may locally alter NE tension and NPC dilation. Consistent with this possibility, there is evidence that mechanical strain on the nuclear lamina is higher at the basal versus apical surface of the nucleus (Carley et al., 2021; Ihalainen et al., 2015). Whether this asymmetry in lamina tension is reflected in the dilatory state of NPCs is unknown. Answering these questions will require facile and accessible methods to examine NPC diameter, ideally at single NPC resolution, in multiple cell types and tissues.

Similar methods are also required to tackle the question of whether NPCs are compositionally (and functionally) different among cell types in both physiological and pathological settings (Cho and Hetzer, 2020; Fernandez-Martinez and Rout, 2021). For example, it has been reported that there are at least two forms of NPCs in budding yeast with either one or two NRs (Akey et al., 2022). Moreover, the nuclear basket is absent from many yeast NPCs and may be assembled as part of a dynamic mRNA export platform (Bensidoun et al., 2022; Galy et al., 2004; Singh et al., 2024). Whether there is such plasticity of the nuclear basket in human cells remains unknown, but there are many hints that there is compositional heterogeneity of NPCs in certain cell types (D’Angelo et al., 2012; Ori et al., 2013) driven by differential nup expression and/or turnover rates, both of which may be influenced by age and disease (Cho and Hetzer, 2020). Importantly, although cryo-ET may be amenable to uncovering broad classes of NPC structures in scenarios where there is a total absence of a given nup (Taniguchi et al., 2024), it will be less sensitive to changes in the relative stoichiometry of nups within individual NPCs.

Another motivation for developing methods to visualize compositional heterogeneity at the individual NPC level is exemplified by evidence that NPCs may be compromised in neurodegenerative disease (Chandra and Lusk, 2022). Specifically, an amyotrophic lateral sclerosis (ALS) pathomechanism caused by a hexanucleotide repeat expansion in the *C9orf72* gene has been proposed to occur through an NPC injury cascade resulting in the loss of a specific subset of nups from a fraction of NPCs (Coyne et al., 2020). POM121, one of three transmembrane nups, is a key linchpin whose disappearance precedes that of other nups, ultimately heralding a loss of nuclear compartmentalization (Baskerville et al., 2024; Coyne et al., 2021). The underlying mechanisms that drive these changes to NPCs remains uncertain but would benefit from a methodology that can reveal the molecular and morphological changes that occur within individual NPCs along the NPC injury cascade. Moreover, the ideal method would couple quantitative immunolabeling with the capacity to assess gross changes in NPC architecture.

Here, we explore the utility of pan-Expansion Microscopy (pan-ExM) to robustly visualize the molecular composition and structure of individual NPCs. Pan-ExM is unique over other expansion microscopy methods as it preserves the total proteome, which can be visualized at an ultrastructural resolution using a “pan” fluorescent protein stain (M’Saad and Bewersdorf, 2020). We demonstrate that machine-learning based segmentation of pan-ExM images robustly and comprehensively identifies individual NPCs, reveals their compositional heterogeneity and allows measurements of their diameter. For example, we recapitulate the observed constriction of NPCs that occurs upon hyperosmotic shock (Zimmerli et al., 2021) while uncovering that NPCs are similarly constricted in annulate lamellae (AL). More strikingly, less dramatic but nonetheless significant diameter differences are observed in NPCs along the basal surface of the nuclear envelope in local “tension islands” that are dependent on LINC complexes. Lastly, we use pan-ExM to discover that in induced pluripotent stem cell derived neurons (iPSNs) from a patient with the pathological *C9orf72* repeat expansion, there is a shift in the distribution of POM121 from the NR to the IR. These data suggest that previously unappreciated, discrete changes to NPC architecture in the *C9orf72* ALS NPC injury cascade are visible by pan-ExM. Thus pan-ExM is a valuable discovery tool that will help illuminate NPC plasticity – an essential step towards understanding its function across cells and tissues.

## Results

### Pan-ExM permits visualization of whole cell proteinaceous ultrastructure including NPCs

To visualize cellular ultrastructure and NPCs, we leveraged the pan-ExM protocol (M’Saad and Bewersdorf, 2020) and fixed and embedded samples in a series of swellable hydrogels, enabling ∼16-fold expansion. A schematic of the procedures and computational tools used in this study are diagrammed in Supplementary Figure Briefly, to “pan” stain all cellular proteins, lysines across the proteome were labeled with a fluorophore conjugated N-hydroxysuccinimide (NHS), DNA was labeled with SYTOX Green, and proteins of interest were labeled with antibodies (Supplementary Figure 1a). Using confocal microscopy, we acquired images capturing whole cell volumes. As expected (M’Saad and Bewersdorf, 2020), cellular organelles like nucleoli, mitochondria, Golgi and centrioles were identifiable by their characteristic ultrastructural morphologies revealed by the pan-stain (Supplementary Figure 1b). NPCs were likewise visible at the edge of the nucleus (Supplementary Figure 1b). We developed an image analysis pipeline using Imaris software and the Fiji (Schindelin et al., 2012) plugin LABKIT (Arzt et al., 2022), a machine-learning based random forest classifier, to segment cellular structures in 3D (Supplementary Figure 1c). Through manual labeling of image voxels in all acquired channels, nuclei were segmented by iterative training. NPCs and component CR, IR and NRs were robustly identified in all axial orientations and automatically segmented by training multiple datasets. This analysis was further combined with segmentation of antibody labeling and AL. Together, these in silico segmentation approaches enabled a detailed ultrastructural analysis of all NPCs in a given cell nucleus in virtually any cell line. Initially, we examined three commonly used cell lines: HeLa cervical adenocarcinoma, A549 lung adenocarcinoma and SH-SY5Y neuroblastoma cells.

In axial sections of magnified views of the nuclear surface, each NPC (regardless of cell line) was comprised of a single focus suspended between two rings (Figure 1a). The former is best observed in cross section (Figure 1b). Consistent with a body of prior work (Krull et al., 2010; M’Saad and Bewersdorf, 2020; Ou et al., 2017; Schermelleh et al., 2008) supporting that NPCs engage euchromatin and/or exclude heterochromatin, the NRs were nestled within islands lacking detectable SYTOX fluorescence (Figure 1a, bottom panels). A quantitative comparison of the average NPC architecture derived from 101,485 NPCs in pan-ExM samples with those derived from *in cellulo* cryo-ET of human NPCs (Mosalaganti et al., 2022; Schuller et al., 2021) indicates that pan-ExM expands the NPC ∼2 fold more along the nuclear transport axis compared to the radial axis (Figure 1c). Indeed, comparing the relative dimensions observed in the expanded NPCs to the average cryo-ET structure of NPCs in intact DLD-1 cell nuclei (Schuller et al., 2021) supports that pan-ExM maintains a faithful relationship between the diameter of the CR, IR and NRs (Figure 1d, e). Moreover, we can also discern that the IR is more proximal to the NR than the CR, a prominent feature of the DLD-1 NPC model (Schuller et al., 2021)(Figure 1f). However, the dimensions along the nuclear transport axis are exaggerated (Figure 1c). Regardless, NPCs and their substructure are easily visualized by pan-ExM with the relative dimensions of the CR, IR and NRs preserved.

**Figure 1.**
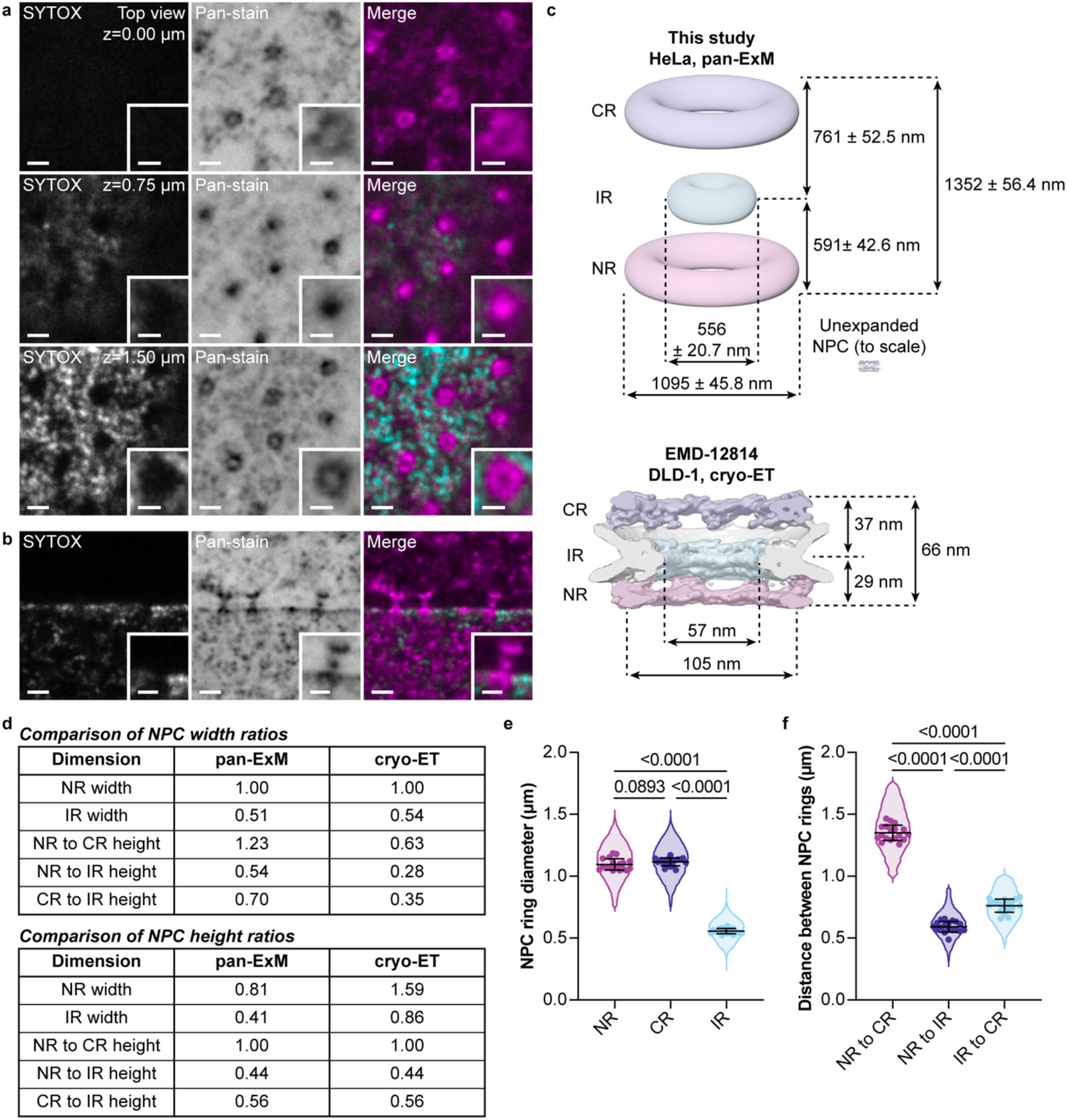
Comparison of pan-ExM expanded and *in cellulo* NPCs a, **b.** Representative confocal fluorescence microscopy images of NPCs in expanded HeLa cells stained with SYTOX and NHS ester pan-stain at indicated axial positions (a) and in cross-section (b). Scale bars 2 µm. Insets show magnified view of an NPC, scale bars 1 µm. **c.** Average NPC dimensions in pan-ExM samples compared to an *in cellulo* cryo-ET NPC density map (Schuller et al., 2021). CR: cytoplasmic ring, IR: inner ring, NR: nuclear ring. **d.** Summary of NPC width and height ratio comparisons in pan-ExM and cryo-ET samples. **e.** Diameter of the indicated NPC rings in pan-ExM expanded HeLa cells. n=101,485 NPC ring diameters total in n=18 cells from a single expansion experiment. Symbols denote individual cell means. Line and error bars are the overall mean ± s.d. Repeated measures one-way ANOVA with Tukey’s multiple comparisons test. **f.** Distance between NPC rings in pan-ExM expanded HeLa cells. n=38,048 NPC ring distances total in n=18 cells from a single expansion experiment. Symbols denote individual cell means. Line and error bars are the overall mean ± s.d. Repeated measures one-way ANOVA with Tukey’s multiple comparisons test.

The ability to segment NPCs and nuclear contours provided a facile approach to automate counting all the NPCs of hundreds of HeLa, A549 and SH-SY5Y nuclei and to correlate these values to nuclear surface and volume measurements. The latter were normalized to an expansion factor calculated for each sample by measuring the mean distance between mitochondria cristae (Supplementary Figure 2a) and centriole diameters (Supplementary Figure 2b); the average of these two was then taken as the expansion factor for assigning pre-expansion scale (Supplementary Figure 2c). There was remarkable consistency in sample-to-sample expansion that ranged from 13.94 to 16.69-fold (Supplementary Figure 2d). On average, the three cell lines had a similar mean number of NPCs with considerable variability on an individual cell basis. HeLa cells displayed the largest spread in values (over 7-fold) and SH-SY5Y cells, the least (Figure 2a). As such, SH-SY5Y cells also had the highest density of NPCs that were closest to each other (Figure 2b-d). To gain some insight into the mechanisms that may influence NPC number, we further related NPC number to the nuclear surface area and nuclear volume on an individual cell basis. Across all cells examined, NPC number was correlated with nuclear surface area (Figure 2e-g) and nuclear volume (Figure 2h-j), with modestly stronger correlations with nuclear volume. Thus, pan-ExM is a valuable and facile approach to count NPCs in relation to nuclear metrics.

**Figure 2.**
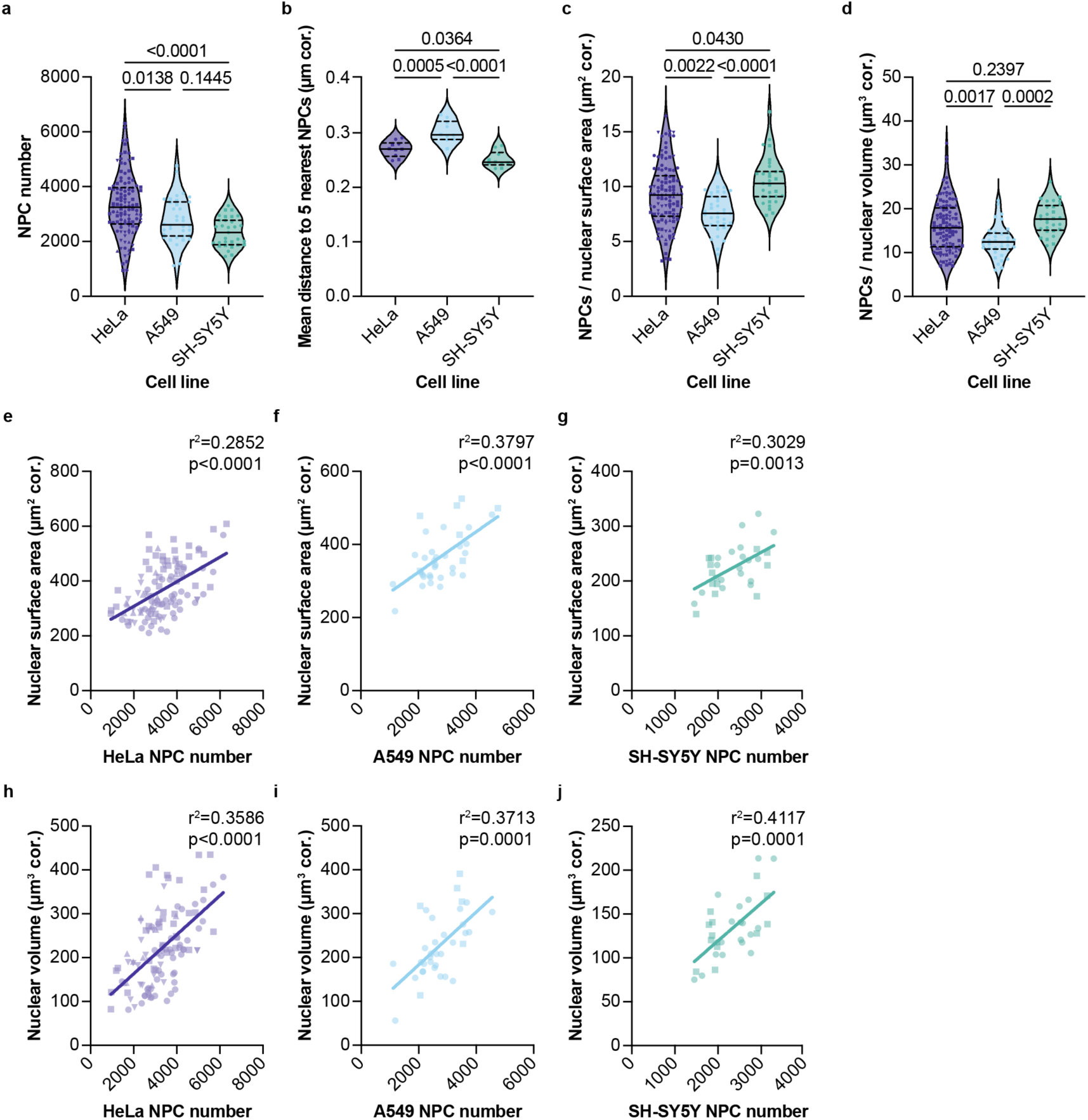
Comprehensive visualization of NPCs with pan-ExM reveals characteristic NPC density and distribution across representative cell lines. **a.** Total NPC number is variable and highest in expanded HeLa cells compared to A549 and SH-SY5Y cells. n=12, n=19, n=33 and n=47 HeLa cells from n=4 independent expansion experiments. n=7 and n=29 A549 cells from n=2 independent expansion experiments. n=11 and n=20 SH-SY5Y cells from n=2 independent expansion experiments. Symbols denote individual cell values and symbol shape indicates expansion experiment. Median values shown as solid lines and quartile values shown as dashed lines. Ordinary one-way ANOVA with Tukey’s multiple comparisons test. **b.** NPCs are most clustered in SH-SY5Y cells. Average distance to the five nearest NPCs (corrected “cor.” for the determined expansion factor) measured in cells representative of local NPC density in each cell line. n=5 cells per line from n=2 independent expansion experiments per cell line. Plot details and statistical tests as in (a). **c.** Overall NPC density (NPCs per nuclear surface area corrected for the determined expansion factor) is variable across the population but highest in SH-SY5Y cells. Plot details and statistical tests as in (a). **d.** Total NPC number per nuclear volume (corrected for the determined expansion factor) is more characteristic than NPC density on the nuclear surface. Plot details and statistical tests as in (a). **e-g.** Total NPC number trends with nuclear surface area but with substantial variability. Symbols as in (a). Lines represent a simple linear regression with the coefficient of determination (r^2^) indicated. **h-j.** NPC number generally correlates better with nuclear volume than nuclear surface area (compare e to h). Symbols as in (a) and regression analysis as in (e-g).

### Nup antibody labeling defines NPC ultrastructure in pan-ExM

We next assessed whether individual nups could be localized within the pan-ExM ultrastructure by immunostaining of expanded cells with a battery of nup specific antibodies. We tested antibodies directed against nups representative of all major NPC architectural elements including the CR, IR and NR, the FG-rich central channel, the nuclear basket, and the cytoplasmic filaments (Figure 3a). The anti-nup antibodies specifically labeled one or more of the three substructures of the expanded NPCs in a manner congruent with their established locations (Figure 3b, c, Supplementary Figure 3a, b). For example, we observed anti-NUP107 and anti-NUP96 staining at both pan-stained rings, confirming these to be the CR and NRs (Supplementary Figure 3c, d). By contrast, the pan-stained focus suspended between the two rings was recognized by antibodies to the central channel FG-nups NUP62 and NUP98, the IR component, NUP93, and the integral membrane nucleoporins NDC1 and GP210. Thus, this middle focus represents the IR with FG-network. Last, although the position of the nuclear basket (NUP50, NUP153, TPR) and cytoplasmic filament (NUP358) labels also converged qualitatively on the pan-stained CR and NRs, quantification of the label position relative to the surface of the nucleus (after normalization to the position of NUP107 to define the “middle” of the NPC) supports that they do, as expected, extend away from the rings into the nucleoplasm and cytoplasm, respectively (Figure 3d). Thus, the overall position of nups within the NPC architecture is retained during fixation and expansion of NPCs.

**Figure 3.**
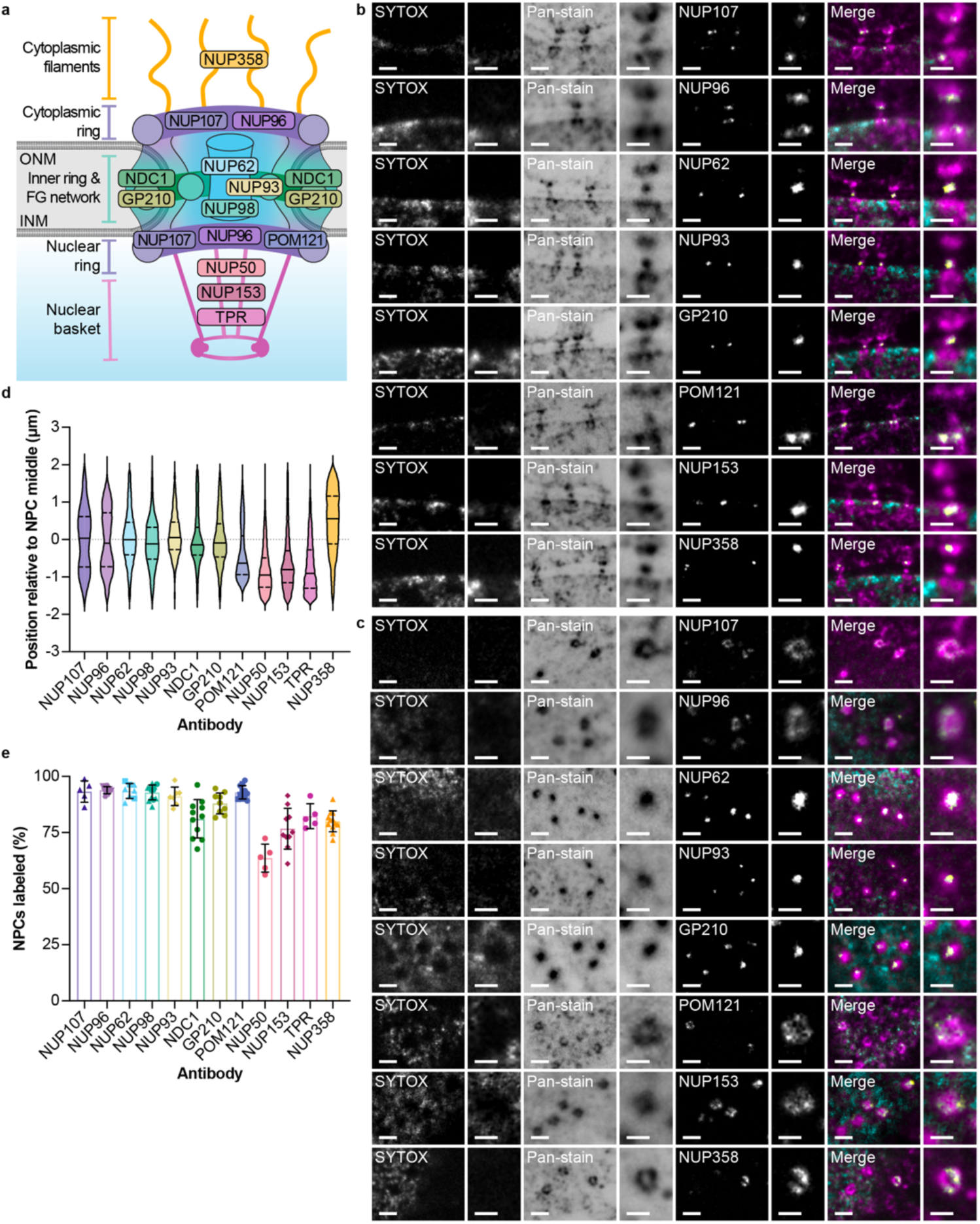
Nup antibody labeling establishes the ability of pan-ExM to reveal nup position within the NPC ultrastructure. **a.** Schematic of the NPC with established nup positions and NPC architectural subunits indicated. ONM: outer nuclear membrane, INM: inner nuclear membrane. **b**, **c.** Representative confocal fluorescence microscopy images of NPCs in cross-section (b) or top-down (c) in expanded HeLa or U2OS CRISPR NUP96-mEGFP cells stained with SYTOX, NHS ester pan-stain, and labeled with antibodies against the indicated nups affirms their localization to the expected NPC subunit and uncovers a bias for POM121 at the NR. Scale bars 2 µm. Insets show a magnified view of a NPC, scale bars 1 µm. **d.** Spatial distribution of nup antibodies along the transport axis in cells relative to the NPC middle denoted by the dotted line reinforces the ability of pan-ExM to faithfully retain established NPC architecture. Median values shown as solid lines and quartile values shown as dashed lines. n=88,581 segmented antibody spots total in n=5 NUP107 labeled HeLa cells from n=1 expansion experiment, n=5 NUP96 labeled U2OS CRISPR NUP96-mEGFP cells each from n=2 independent expansion experiments, n=3 and n=5 NUP62 labeled HeLa cells from n=2 independent expansion experiments, n=3 and n=9 NUP98 labeled HeLa cells from n=2 independent expansion experiments, n=7 NUP93 labeled HeLa cells from n=1 expansion experiment, n=11 NDC1 labeled HeLa cells from n=1 expansion experiment, n=9 GP210 labeled HeLa cells from n=1 expansion experiment, n=5 and n=6 POM121 labeled HeLa cells from n=2 independent expansion experiments, n=5 NUP50 labeled HeLa cells from n=1 expansion experiment, n=10 NUP153 labeled HeLa cells from n=1 expansion experiment, n=5 TPR labeled HeLa cells from n=1 expansion experiment and n=12 NUP358 labeled HeLa cells from n=1 expansion experiment. Median values shown as solid lines and quartile values shown as dashed lines. **e.** Nearly all NPCs segmented based on the NHS ester-pan stain were labeled with antibodies to the CR, IR and NR components whereas the peripheral nups were detected at most but not all NPCs. Symbols denote individual cell values and symbol shape indicates expansion experiment. Bars and error bars are the mean ± s.d.

Because the overall structure of NPCs and the relative position of all tested nups was preserved in the pan-stain, we more closely evaluated the localization of the transmembrane nup, POM121. Indeed, although a common theme across eukaryotes is that the transmembrane nucleoporins engage the IR of the NPC, biochemical (Yavuz et al., 2010) and cross-linking mass spectrometry (Bui et al., 2013) evidence support that it may engage the CR and NR nups as well, perhaps in a mutually exclusive manner (Mitchell et al., 2010). Interestingly, unlike antibodies directed to the other transmembrane domain-containing nups, GP210 and NDC1, which stained the IR as expected (Figure 3), POM121 antibodies exclusively labeled the nuclear aspect of the NPC most closely mimicking the positions of NUP107 and NUP96 at the NR (Figure 3d). This result was consistent with a recent super-resolution microscopy study (using a different antibody)(Andronov et al., 2022) and was not unique to HeLa NPCs, as we observed the identical asymmetry in the POM121 labeling of A549 and SH-SY5Y cells as well (Supplementary Figure 3e, f). Thus, POM121 is unique amongst the transmembrane nups and is asymmetrically distributed in the NPC on the NR.

As the pan-stain identifies all NPCs and major architectural units, a major advantage of this approach is the ability to directly assess antibody labeling efficiency – a necessity to unambiguously detect potential changes in nup composition at the individual NPC level. For the central channel FG-nups and the scaffold elements of the NPC including the CR, IR and NRs, we observed specific and near-comprehensive labeling of the pan-stained NPCs (Figure 3e). In contrast, the labeling of the transmembrane nup NDC1, the nuclear basket nups NUP50, TPR and NUP153 and the cytoplasmic filament nup Nup358 was notably less efficient (64-82%). Thus, it is possible there are sub populations of NPCs that lack these asymmetric elements, in line with observations in budding yeast (Akey et al., 2022; Bensidoun et al., 2022; Galy et al., 2004; Niepel et al., 2013; Singh et al., 2024). Despite the overall labeling efficiency of NPCs, we note that there is likely the loss of some nup epitopes during the expansion procedure, explaining why we were unable to always visualize a ring-like staining pattern. Consistent with this, although recent work indicates that some molecules of NUP93 and NUP62 reside at CRs (Bley et al., 2022; Huang et al., 2020; Mosalaganti et al., 2022; Zhu et al., 2022), these pools are only rarely observed in the immunolabeling of the pan-stained samples (Supplementary Figure 3g).

### Local differences in NPC dimensions

Emerging work has intimated that the NPC diameter adopts a range of dilation states in response to NE tension, which in turn may be modulated by both cell-intrinsic and extrinsic factors (Mosalaganti et al., 2022; Schuller et al., 2021; Zimmerli et al., 2021). Although an exciting concept, we know little about the factors that govern if NPCs in a single nucleus exist in different dilatory states and, if so, if this is biologically meaningful. While the most pronounced quantitative changes in NPC diameter when comparing a no tension state (isolated NEs) and a physiological tension state (*in cellulo*) manifest at the IR, which expands the central channel ∼30% in two different cryo-ET models of human cells (Mosalaganti et al., 2022; Schuller et al., 2021), changes in tension are also reflected in the pore membrane to pore membrane distance (a 12-18% expansion in human cells) (Mosalaganti et al., 2022; Schuller et al., 2021). The effect of NE tension on the dilation of the CR and NRs is more modest and most clearly revealed by comparing fission yeast NPC cryo-ET structures in different NE tension states – here a 13% change in the NR diameter was observed (Zimmerli et al., 2021). It is important to note, however, that the diameter of individual NPCs in the *in cellulo* cryo-ET models varies considerably (Akey et al., 2022; Mahamid et al., 2016; Schuller et al., 2021; Zimmerli et al., 2021). To test whether pan-ExM could provide a tool to readily evaluate relative NPC diameter within individual cells at high sampling density, we measured the diameters of thousands of CR, IR and NRs of computationally segmented NPCs from several cells in multiple independently expanded samples. In line with prior observations (Schuller et al., 2021), we noted that there was a near two-fold range in NPC diameter values across the nuclei (Figure 4a). We investigated if the diameter variance reflected any biases in the distribution of NPCs across the nuclear surface. As there is evidence that tension on the nuclear lamina is higher on the bottom of the nucleus due to cell adhesion to the extracellular matrix (Carley et al., 2021; Ihalainen et al., 2015), we first contrasted NPC ring diameters on the top and bottom of the nucleus. Consistent with the expectation that higher force transduction to the nuclear lamina increases NE tension in a way that could enhance NPC dilation, we observed a significant trend of larger NPC ring diameters on the basal surface compared to the apical surface (Figure 4a) although there was marked variance in diameters on both surfaces. One interpretation of these data is that there are localized regions of high NE tension, particularly on the basal nuclear surface.

**Figure 4.**
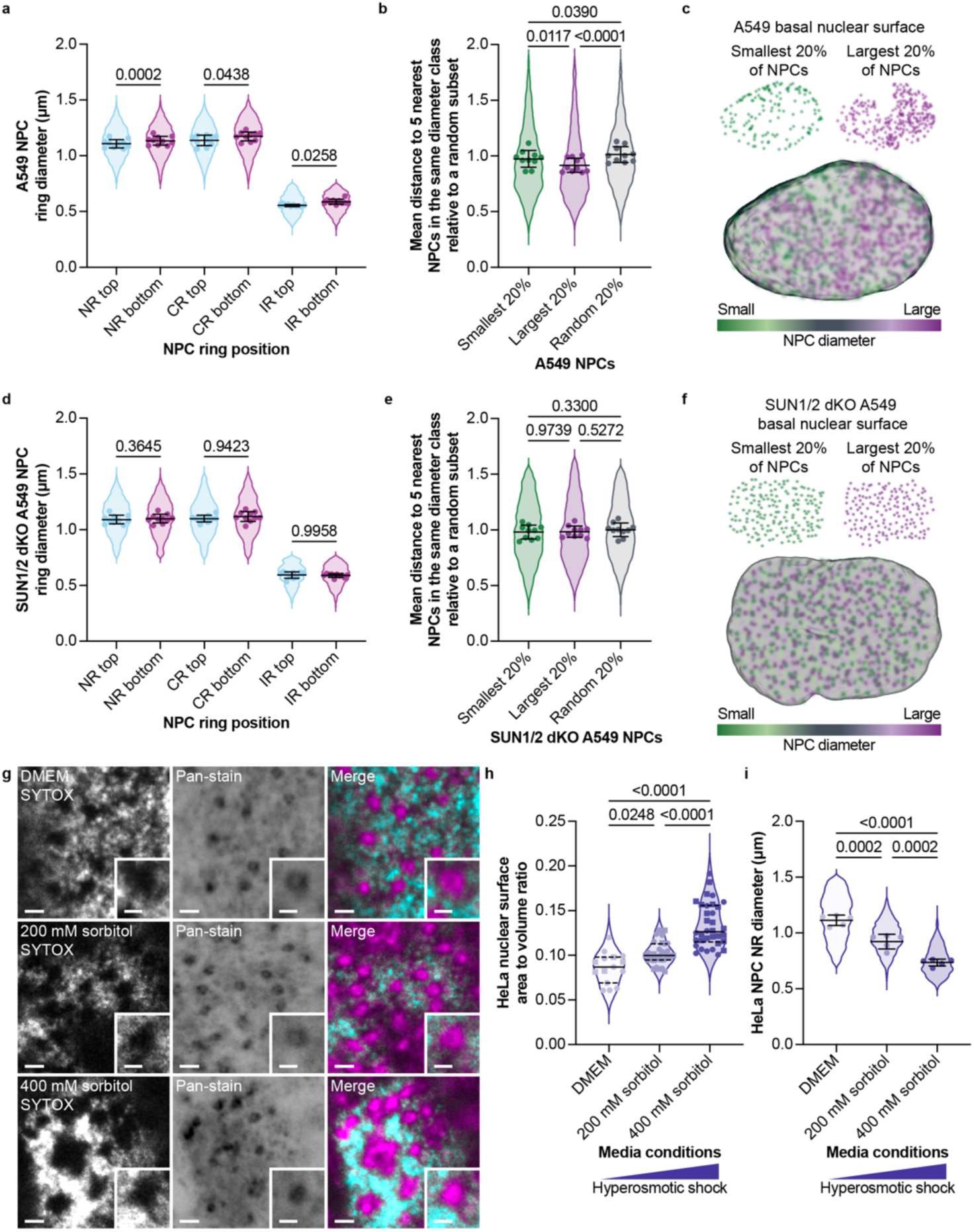
Pan-ExM reveals LINC complex-dependent, local differences in NPC diameter and NPC constriction in hyperosmotic conditions. **a.** The nuclear ring (NR), cytoplasmic ring (CR) and inner ring (IR) of NPCs at the bottom (closest to the basal cell surface) of the nucleus are dilated compared to those at the top in A549 cells. n=21,593 NPC rings total in n=10 cells from n=1 expansion experiment. Symbols denote individual cell means. Median values shown as solid lines and quartile values shown as dashed lines. Repeated measures one-way ANOVA with Tukey’s multiple comparisons test. Independent expansion included in Supplementary Figure 4a. **b.** NPCs are more likely to reside near neighboring NPCs of like diameter. Average distance to the five nearest NPCs within the same diameter class relative to a random subset of NPCs from the same nucleus. n=10,846 NRs total in n=10 cells from n=1 expansion experiment. Plot details and statistical analysis as in (a). Independent expansion included in Supplementary Figure 4b. **c.** Visualization of NPC distribution according to dilation state at the basal nuclear surface of an A549 cell by color-coding according to NPC diameter class. The most constricted 20% of NPCs are shown in dark green, and the most dilated 20% are shown in dark purple. All NPCs are overlaid on a 3D rendering of the nucleus. **d.** Disrupting LINC complexes by CRISPR ablation of *Sun1* and *Sun2* (SUN1/2 dKO) leads to a loss of the bias for greater NPC dilation on the bottom of the nucleus. n=14,627 NPC rings total in n=10 cells from n=1 expansion experiment. Plot details and statistical analysis as in (a). Independent expansion included in Supplementary Figure 4l. **e, f.** Disrupting LINC complexes leads to homogenization of NPC diameter across the nuclear surface. n=9,475 NRs total in n=10 cells from n=1 expansion experiment. Plot details and statistical analysis as in (a). Independent expansion included in Supplementary Figure 4m. **g**. Representative confocal fluorescence microscopy images of NPCs in expanded HeLa cells in isotonic (DMEM) or hyperosmotic media (sorbitol) with SYTOX and NHS ester pan-stain. Scale bars 2 µm. Insets show magnified view of a NPC, scale bars 1 µm. **h.** NE tension is reduced in hyperosomotic shock conditions as quantified by an increase in the nuclear surface area to volume ratio. n=5 and n=10 cells in DMEM, n=15 cells in 200 mM sorbitol each, and n=14 and n=20 cells in 400 mM sorbitol from n=2 independent expansion experiments. Symbols denote individual cell values and symbol shape indicates expansion experiment. Median values shown as solid lines and quartile values shown as dashed lines. Ordinary one-way ANOVA with Tukey’s multiple comparisons test. **i.** Hyperosomotic shock leads to NPC constriction. n=5,948 NRs total in n=5 cells each per media condition from n=1 expansion experiment. Plot details and statistical analysis as in (h).

Interestingly, upon binning the NPC diameters into quintiles, the distance to the five nearest NPCs in the same diameter class was shortest for NPCs with the largest 20% of diameters (Figure 4b). Indeed, compared to a randomly selected 20%, the largest NPCs were significantly closer together. Color-coding of NPCs at the basal nuclear surface according to NPC diameter class revealed that larger NPCs locally clustered in apparent hot spots (Figure 4c, purple). These results were consistently reproduced in an independent expansion experiment and mirrored in HeLa cells (Supplementary Figure 4a-d). Thus, the largest NPCs cluster together in local regions across the nuclear surface.

To investigate whether the measured biases in NPC diameter reflected the functional integration of NPCs within a mechanoresponsive network (and to rule out expansion-induced artifacts), we performed pan-ExM and measured NPC diameters in CRISPR-edited A549 cells lacking *Sun1* and *Sun2* (Supplementary Figure 4e-k). SUN1 and SUN2 are integral components of LINC complexes (King, 2023) necessary to transmit cytoskeletal forces from cell-matrix adhesions to the nuclear lamina (Carley et al., 2021). Strikingly, the ablation of LINC complexes completely disrupted the observed bias for larger NPC diameters on the basal surface of the nucleus in cell populations from two independent expansion experiments (Figure 4d, Supplementary Figure 4l), which were now also found to be uniformly distributed. Further, we no longer detected a bias for the largest NPCs to cluster (Figure 4e, f, Supplementary Figure 4m). Thus, pan-ExM helped detect LINC-complex dependent “tension islands” where NPC diameter was locally more uniform suggesting a model in which LINC complexes directly or indirectly modulate the diameter of subsets of NPCs.

As we observed that LINC complex ablation had a relatively modest quantitative effect on NPC diameter, we wondered whether this perturbation reflected the complete relaxation of NPCs to their most constricted state. As hyperosmotic shock is thought to constrict NPCs (Zimmerli et al., 2021), we tested how NPC diameter was impacted by treating cells with high concentrations of sorbitol. As shown in Figure 4g-i, treatment of HeLa cells with increasing concentrations of sorbitol led to a marked increase in the surface area to volume ratio of nuclei and a commensurate decrease in NPC diameter (as measured at the NR). Note that the relative change of NR diameter from the osmotically equilibrated state (DMEM) to those in 400 mM sorbitol was ∼40%. In contrast, the change between the WT and SUN1/2 dKOs A549 cells was ∼5% (Supplementary Figure 4n). Taken together, these observations suggest that the embedding of NPCs in the taut NE membrane drives the majority of the observed dilation while the extent of orthogonal tension exerted by LINC complexes contributes more modestly to NPC dilation/constriction.

### Pan-ExM reveals organization of NPCs in AL

We next assessed whether pan-ExM could be used to provide insight into the dilatory state, molecular composition, and 3D organization of AL, which were robustly observed in an iPSC line and were occasionally present in HeLa and SH-SY5Y cells. As expected from prior EM studies (Cordes et al., 1996; Kessel, 1983), the NPCs in AL were organized in densely stacked arrays that were present in both an en face (top-down; Figure 5a) and lateral (Figure 5b) orientations. As the AL would not be predicted to be under tension, NPC diameter in AL was markedly (∼35%) reduced compared to NPCs at the NE (Figure 5c, d) and in line with the hyperosmotic treatment. Thus, it is likely that NPCs in AL are found in a fully constricted state.

**Figure 5.**
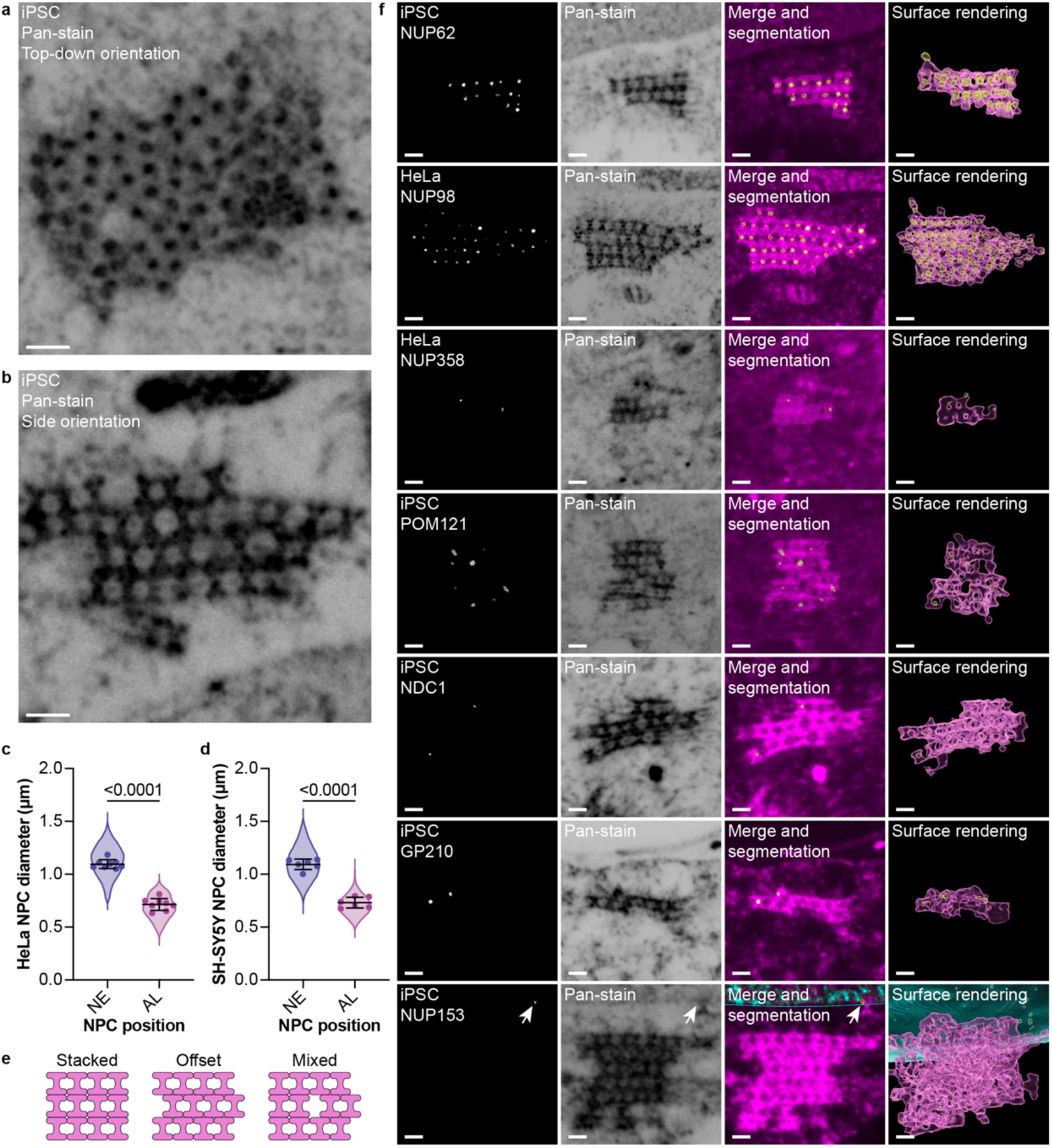
Pan-ExM reveals organization of annulate lamellae a, **b.** Stacks of AL – NPCs embedded in the endoplasmic reticulum – can be readily identified by their ultrastructure in pan-ExM samples. Representative confocal fluorescence microscopy images of AL of expanded iPSCs stained with NHS ester pan-stain in top-down (a) and side view (b) orientations. **c, d.** NE-embedded NPCs are dilated compared to those in AL in HeLa and SH-SY5Y cells. n=11,606 NE NPC NRs and AL NPCs total in n=8 HeLa cells from n=1 expansion experiment. n=1,821 NE NPC NRs and AL NPCs total in n=6 SH-SY5Y cells from n=1 expansion experiment. Symbols denote individual cell means. Line and error bars are the overall mean ± s.d. Paired t-test. **e.** The organization of individual NPCs in AL ranges from stacked, to offset, or a mixture as illustrated in the schematics. **f.** NPCs in AL are readily stained with antibodies to the FG-nups but are sparsely labeled with antibodies to the transmembrane nups or asymmetric NPC elements. Representative confocal fluorescence microscopy images of AL in expanded iPSC and HeLa cells labeled with antibodies against the indicated nups, stained with NHS ester pan-stain. 3D surface renderings of segmented structures are also shown. Arrow points to NUP153 labeling of an NPC embedded in the NE. Scale bars 2 µm.

While EM favors efficient labeling of membranes, we expected that the total protein labeling in pan-ExM could provide new insight into putative interactions between NPCs that may underlie the biogenesis of AL. Indeed, the outer rings of the NPCs appeared to directly connect to NPCs both above and below in two conformations (Figure 5e). In one, the NPCs were stacked on top of each other such that the transport channels aligned. In another, the NPCs were offset in a running bond brick-like pattern. Most AL comprised a mixture of these two arrangements. In principle, the observed juxtapositions of NPCs could be driven by interactions between asymmetric elements of NPCs i.e. the cytoplasmic filaments and nuclear basket, or between symmetric elements (i.e. the outer rings).

To test these models of NPC stacking in AL, we immunolabeled cells with nup antibodies (Figure 5f). Whereas antibodies directed towards the FG-nups, NUP62 and NUP98, robustly labeled AL, we only observed sparse labeling of the cytoplasmic filament nup NUP358 and, surprisingly, the transmembrane nups POM121, NDC1 and GP210. We were unable to identify the basket component NUP153 in AL stacks, despite clear NUP153 staining at NPCs embedded in the NE in the same expanded cell (Figure 5f, arrow). Thus, our observations, when taken together with other studies on *Xenopus* (Rasala et al., 2008; Walther et al., 2003) and proteomic analyses and cryo-ET of AL from *Drosophila* embryos (Hampoelz et al., 2016; Hampoelz et al., 2019; Sachweh et al., 2024) support that NPCs in AL have reduced levels of transmembrane nups and lack the asymmetric elements of the NPC. Together, we favor the hypothesis that direct outer ring interactions facilitate the NPC stacking underlying AL formation. However, we are cognizant that NPCs “over expand” along the transport axis, which could also influence the observed arrangement of NPCs in AL.

### POM121 shifts position in model ALS iPSNs

To further push the boundaries of pan-ExM and to investigate the potential functional significance of POM121’s specific localization to the NR, we tested whether pan-ExM could distinguish compositionally unique NPCs in the context of a neurodegenerative disease model. Recent work supports that there is an NPC injury cascade in which POM121 plays a critical role in the characteristic loss of a subset of nups from NEs in iPSC derived neurons (iPSNs) from patients expressing a hexanucleotide repeat expansion in *C9orf72,* the most common genetic cause of ALS (Coyne et al., 2020). The observed NPC injury occurs approximately 32 days after differentiation into motor neurons in culture. We therefore prepared differentiated iPSNs at day 18 (before NPC injury) and day 32 (after NPC injury) post-differentiation for examination by pan-ExM. In all samples, NPCs were readily visible in the pan-stain (Figure 6a). Notably, while the extent of immunolabeling of NUP62 (previously found to be unaffected in this disease model (Coyne et al., 2021; Coyne et al., 2020) showed no change between day 18 and day 32 iPSNs (Supplementary Figure 5), we observed a marked reduction in the labeling of POM121 (Figure 6b) from 92% of NPCs at day 18 to 72% at day 32 in *C9orf72* HRE-expressing iPSNs. POM121 labeling of NPCs was also modestly reduced in control iPSNs from 91% of NPCs at day 18 to 86% of NPCs at day 32. Prior studies have suggested that loss of POM121 is a hallmark of a broader disruption in NPC composition including diminished levels of seven other nups across multiple subcomplexes (Coyne et al., 2020). Consistent with this, we also observed that NPCs lacking POM121 have a statistically significant decrease in pan-staining intensity within the segmented NPCs compared to POM121-containing NPCs in the same nucleus (Figure 6d). Loss of POM121 also correlated with more constricted NR diameters in both control and *C9orf72* HRE-expressing iPSNs at day 18 and day 32 (Figure 6e, f). Thus, pan-ExM allows the visualization of pathological changes to the biochemical identity of individual NPCs.

**Figure 6.**
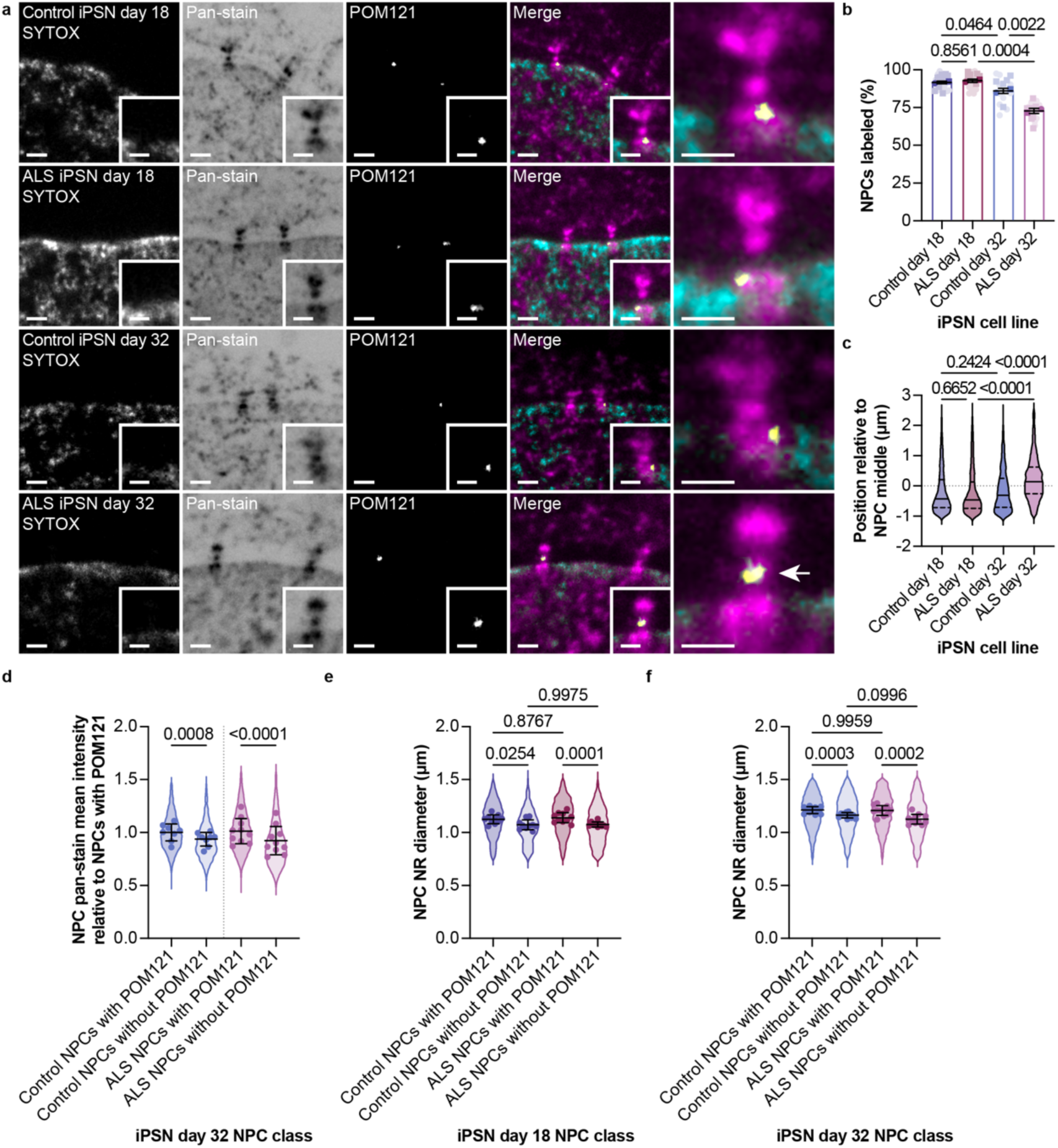
POM121 shifts to the inner ring and is ultimately lost from NPCs in aged model ALS iPSNs. **a.** POM121 is detected at the IR rather than the NR in “aged” model ALS iPSNs after 32 days of differentiation. Representative confocal fluorescence microscopy images of NPCs in expanded iPSNs derived from a patient with ALS or from an age and sex-matched healthy control donor at indicated number of days after differentiation stained with SYTOX, NHS ester pan-stain, and labeled with an antibody against POM121. Scale bars 2 µm. Insets and far right panel show magnified views of a NPC, scale bars 1 µm. Arrow points to labeling of the IR. **b.** The fraction of NPCs labeled with POM121 antibody declines subtly over days of differentiation in control iPSNs but dramatically in ALS patient-derived iPSNs. Percentage of NPCs per nucleilabeled with an antibody against POM121 in iPSNs of the indicated genotype and number of days after differentiation. n=11 and n=13 control day 18 iPSNs, n=11 and n=9 ALS day 18 iPSNs, n=9 control day 32 iPSNs each, and n=11 and n=10 ALS day 32 iPSNs from n=2 independent expansion experiments. Symbols denote individual cell means and the independent expansion experiments means (darker symbols). Bars and error bars are the overall mean ± s.d. Ordinary one-way ANOVA with Tukey’s multiple comparisons test. **c.** Antibody staining of POM121 is shifted from the NR to the IR across the population of POM121-positive NPCs specifically in “aged” model ALS iPSNs. Spatial distribution of the indicated genotype and number of days after differentiation relative to NPC middle denoted by the dotted line. n=8,377 segmented POM121 antibody spots total in n=5 cells per cell line each from n=2 independent expansion experiments. Median values shown as solid lines and quartile values shown as dashed lines. Kruskal-Wallis test with Dunn’s multiple comparisons test. **d.** NPCs lacking POM121 staining have reduced overall pan-stain signal compared to POM121-positive NPCs in the same nucleus. Quantification of pan-stain mean intensity at segmented NPCs relative to segmented NPCs with POM121 in iPSNs of the indicated genotype at day 32 post differentiation. n=22,427 segmented NPCs total in n=8 control day 32 iPSNs and n=10 ALS day 32 iPSNs from n=1 expansion experiment. Symbols denote individual cell means. Line and error bars are the overall mean ± s.d. Paired t-tests. **e, f.** ALS iPSN NPCs lacking POM121 staining are more constricted than POM121-positive NPCs in the same nucleus at both day 18 (e) and day 32 (f) post differentiation. n=12,560 NRs total in n=10 day 18 iPSNs each from n=1 expansion experiment. n=13,323 NRs total in n=11 day 32 iPSNs each from n=1 expansion experiment. Symbols denote individual cell means. Line and error bars are the overall mean ± s.d. Paired t-tests or ordinary one-way ANOVA with Tukey’s multiple comparisons test for comparing NPCs within the same cell or between cells, respectively.

Most strikingly, upon close inspection of the anti-POM121 staining in the day 32 *C9orf72* HRE-expressing iPSNs, it was clear that in NPCs that retained POM121 it was no longer distributed along the NR (Figure 6a, bottom right panel). Indeed, in virtually all the antibody-labeled NPCs, the POM121 stain was now re-positioned instead to the IR while the anti-POM121 labeling in the iPSN controls at the same timepoint retained the normal NR localization (Figure 6a, c). Thus, using pan-ExM we have visualized a remarkable response in the position of a key transmembrane nup from the NR to the IR in a pathological condition. While the underlying mechanism driving these changes remains uncertain, we suggest that this may be an important, but to this point invisible, step along the NPC injury cascade that may contribute to an ALS pathomechanism.

## Discussion

We have investigated the ability of pan-ExM to serve as an enabling tool for exploring several pressing questions in the nuclear transport field. We suggest that pan-ExM fills an important niche between cryo-EM/ET and super-resolution microscopy. The pan-ExM approach overcomes the significant financial, technical and intellectual resources required for cryo-EM/ET while also circumventing the modality-specific limitations of super-resolution imaging that are compounded by the need to access specialized microscopes. Pan-ExM particularly excels at providing insight into the molecular composition and structure (at tens of nanometer resolution) at the level of individual NPCs in a given nucleus. As it is possible to perform pan-ExM on virtually any cell type and, with protocol modifications, tissues (M’Saad and Bewersdorf, 2020; M’Saad et al., 2022), it promises to reveal a broad spectrum of NPC diversity that has previously gone underappreciated. The ability to confidently visualize NPCs of unique molecular composition and structure, as demonstrated here, is the first step to understanding their function in both physiological and pathological contexts.

As an example of a pathological context in which pan-ExM was illuminating, we investigated how NPCs change in an iPSN model of *C9orf72* ALS. The data first reinforce that pan-ExM can be used to assess the presence/absence of individual nups from single NPCs, recapitulating the observed loss of POM121 during a *C9orf72* HRE-specific NPC injury cascade thought to be central to an ALS pathomechanism (Coyne and Rothstein, 2021; Coyne et al., 2020). While making a definitive conclusion from negative data (i.e. lack of immunolabeling) is always fraught, pan-ExM has several key advantages over classic immunofluorescence and super-resolution approaches that allows one to arrive at more confident conclusions. First, the pan-stain recognizes virtually all NPCs. This provides a facile and quantitative approach to establish the labeling efficiency per NPC for a given antibody in a manner that is challenging or impossible for other methods. Labeling efficiency is also greatly enhanced by expansion itself, overcoming epitope masking that can occur in crowded macromolecular structures and facilitating efficient penetration of antibodies into the sample (M’Saad and Bewersdorf, 2020). Indeed, we observed that the tested antibodies recognizing scaffold nups label virtually all NPCs, even if the staining patterns suggest some loss of epitopes through the expansion procedure (Figure 3c). Second, pan-ExM revealed a remarkable change to the nanoscale distribution of POM121 that occurred specifically in the *C9orf72* HRE-expressing iPSN line 32 days after differentiation into motor neurons (Figure 6). The timing is remarkable: precisely when NPC injury has been described to begin, we observe that POM121 shifts position from the NR to the IR. We suspect that this change precedes the complete loss of POM121 from NPCs (Figure 6b), at which time we observe a decrease in the total bulk protein at the NPC (Figure 6c). Thus, pan-ExM revealed a new step in the NPC injury pathway; moreover, this finding suggests that there are changes in biochemical interactions among nups that likely precedes their loss from the NPC, which may also be reflected in the observed constriction of NPCs that lack POM121 (Figure 6d, e). This work thus also reveals an aspect of NPC plasticity that was previously unappreciated.

The underlying mechanism driving the shift in POM121 distribution remains unknown, but as there is evidence that POM121 can biochemically engage with both NUP160 and NUP155 in a mutually exclusive fashion (Mitchell et al., 2010), it is attractive to consider the hypothesis that POM121 moves from a NUP160-bound state (at the NR) to a NUP155-bound state (at the IR). While there are many plausible mechanisms that may contribute to such a change including putative post-translational modifications, another possibility is that POM121’s location reflects its engagement with nuclear transport receptors (NTRs). POM121 is unique among the integral membrane nucleoporins in that it engages directly with the Kap α/Kap β1 NTR complex through a nuclear localization signal in its N-terminus (Rasala et al., 2008; Yavuz et al., 2010). This NLS is thought to be a key functional element that helps to target it to the inner nuclear membrane during early steps of NPC biogenesis (Funakoshi et al., 2011; Talamas and Hetzer, 2011). Interestingly, a previous study indicated that under conditions where POM121 binds Kap α/Kap β1 it displays strong binding to NUP155 but not NUP160 (Yavuz et al., 2010). The release of Kap α binding occurs at the nuclear basket, predominantly through a mechanism requiring NUP50; the NUP50 orthologue in budding yeast, Nup2, is also essential for NLS-dependent targeting of integral membrane proteins to the inner nuclear membrane (King et al., 2006; Lokareddy et al., 2015). Thus, NUP50-dependent release of Kap α from POM121 at the nuclear basket could promote its targeting to the NR. Interestingly, loss of NUP50 appears to be a critical component of NPC injury in ALS (Freibaum et al., 2015; Megat et al., 2023). Thus, it is plausible that the disrupted release of POM121 from Kap α promotes its preference for binding to NUP155 at the IR. This and the ultimate function of POM121 at the NR will be topics of future work that will be supported by the unique ability of pan-ExM to reveal positional information of POM121 and other nups at the nanoscale.

While our discovery of biased positioning of POM121 at the NR of the NPC was entirely unexpected, the ability of pan-ExM to reveal a range of NPC diameters across the same nucleus fills a critical gap in the field. Indeed, approaches are needed to interrogate the contexts and consequences of NE tension on NPC form and function. Changes in NPC diameter have been proposed for decades, but only recently with “*in cellulo*” cryo-ET of NPC structures has definitive evidence supporting this type of NPC plasticity come to light (Akey et al., 2022; Hoffmann et al., 2024; Mahamid et al., 2016; Mosalaganti et al., 2022; Schuller et al., 2021; Taniguchi et al., 2024; Zila et al., 2021; Zimmerli et al., 2021). Here, we provide a more nuanced view of NPC diameter that supports that there is a spectrum of NPC dilatory states with a bias for the most dilated NPCs to be on the basal nuclear surface. The data further suggest that there are local islands (hundreds of square nanometers in dimensions) of NPCs that are more or less dilated. As both the local islands and basal bias of dilated NPCs are abolished upon ablation of LINC complexes, it is most likely that NE tension, driven at least in part by cytoskeletal forces, can in fact modulate NPC diameter. Such an idea may provide a function for the long-observed association of SUN1 with NPCs (Liu et al., 2007; Talamas and Hetzer, 2011), and may suggest a direct mechanoresponsive mechanism to dilate or constrict NPCs. Although the idea that NE tension may impact NPC dilation has been proposed (Elosegui-Artola et al., 2017; Mosalaganti et al., 2022; Schuller et al., 2021; Zimmerli et al., 2021), this work reveals an additional nuance: that this tension may be much more localized than previously thought. It remains mysterious, however, whether these local NE tension differences can manifest actual functional changes to these NPCs in their ability, for example, to establish a selective transport channel and/or impact genome function, particularly as the relative mean changes in NPC diameter appears modest with a change of only ∼5%. Interestingly, recent studies have implicated SUN1 and LINC complex components as contributors to ALS pathophysiology (Baskerville et al., 2024; Sirtori et al., 2024). Specifically, SUN1 may be a critical mediator of passive permeability of the NPC (Baskerville et al., 2024). Although the mechanisms underlying these events remain unknown, it is perhaps not coincidental that loss of POM121 from ALS NPCs correlates with a diameter change of NPCs. In either case, our work suggests that pan-ExM will likely be a useful tool for providing insight into these neurodegenerative events.

More dramatic changes in NPC diameter of ∼35% are observed upon hyperosmotic shock or in the context of NPCs in AL. Collectively these data suggest that the mechanical state of the nucleus more modestly affects NPC dilation than that realized in response to changes in global NE tension, for example that exerted by the osmotic pressure within the nucleus (Hoffmann et al., 2024). Thus, complete constriction of the NPC likely only occurs under extreme environmental perturbation. In these scenarios, it may be logical for cells to attempt to attenuate nuclear transport by “closing” their NPCs with the understanding that the transport channel is far from occluded even in the most closed conformation (Mahamid et al., 2016; Mosalaganti et al., 2022; Schuller et al., 2021; Zimmerli et al., 2021).

The constriction of NPCs in AL also raises the possibility that the integration of NPCs into a more “tense” NE may promote the assembly of NPCs from an immature state. For example, NPCs in AL lack key components including the entirety of the nuclear basket (Hampoelz et al., 2016; Hampoelz et al., 2019; Rasala et al., 2008; Sachweh et al., 2024; Walther et al., 2003). The underlying mechanism explaining why AL NPCs cannot support nuclear basket assembly remains to be defined. One possibility is that, as our data suggest, there are stacking interactions between the outer rings of the NPCs in AL that preclude basket assembly. An alternative model is that basket assembly may only occur in membranes under tension. Such a concept aligns with work exploring the steps in de novo NPC assembly where the basket is curiously added as a terminal step (Onischenko et al., 2020; Otsuka et al., 2023). Further, during post-mitotic NPC assembly, early intermediates are assembled into a small nuclear pore before it dilates to complete assembly (Otsuka et al., 2018). More broadly, this interpretation invites the idea that NE tension could directly impact NPC composition. For example, during zebrafish embryonic development NPCs are thought to mature from a constricted state lacking the nuclear basket to a more mature, transport-competent form after the maternal to zygotic transition (Shen et al., 2022). It also has implications for mechanotransduction mechanisms – for example, the local modulation of tension could favor/disfavor the stabile incorporation of nuclear basket proteins, which could direct RNA export, for example, to biochemically polarized NPCs. Our hope is that pan-ExM will provide a key tool to begin to test these and other hypotheses.

## METHODS

### Cell culture

HeLa cells were cultured in Dulbecco’s modified Eagle medium (DMEM; Gibco, 11965092) supplemented with 10% fetal bovine serum (FBS; Gibco, A5256801), penicillin-streptomycin mix (pen/strep; Gibco, 15140122) and sodium pyruvate (Gibco, 11360070). A549 cells were cultured in DMEM F12 (Gibco, 11320032) with 10% FBS and pen/strep. SH-SY5Y cells were cultured in Eagle’s Minimum Essential Medium (EMEM; ATCC, 30-2003) with 15% heat inactivated FBS, pen/strep and 2 mM Glutamax (Gibco, 35050061). U2OS CRISPR NUP96-mEGFP cells (Cytion, 300174) were cultured in McCoy’s 5A medium (Cytiva, SH30200.01) supplemented with 10% FBS, pen/strep, 2 mM Glutamax, sodium pyruvate and MEM Non-Essential Amino Acids (NEAA; Gibco, 11140050). iPSCs were grown on Geltrex (Thermo Scientific, A1413302) coated plates and cultured in mTESR1 media (Stem Cell Technologies, 85850). All cells were maintained at 37°C with 5% CO_2_. Passaging was performed using 1X PBS and 0.05% Trypsin (Gibco, 25300054) or 0.5 mM EDTA (Corning, 46-034-CI) for iPSCs. 24 h before fixation, ∼65,000 cells were seeded onto coverslips coated with 50 µg/mL collagen (Corning, 354236) or Matrigel (Corning, 356231) for iPSCs.

For hyperosmotic shock treatment, media was aspirated from HeLa cells and replaced with either DMEM media with 400 mM sorbitol, DMEM media with 200 mM sorbitol or DMEM media for 5 min before fixation.

### Direct-induced motor neuron differentiation

*C9orf72* ALS patient and control iPSCs (Supplementary Table 1) were obtained from the Answer ALS repository at Cedars Sinai and differentiated into spinal motor neurons as previously described following a modified direct-induced motor neuron differentiation protocol (Baskerville et al., 2024; Coyne et al., 2020). iPSNs were cryopreserved in Cryostor CS10 media on day 12 of differentiation. Briefly, iPSNs at day 12 of differentiation were thawed and grown on Matrigel (Corning, CLS35623) coated dishes and cultured in stage 3 media composed of 47.5% Iscove’s modified Dulbecco’s medium (IMDM; Gibco, 12440061), 47.5% F12 (Gibco, 11765054) with 2% B-27 (Gibco, 17504044), 1% MEM NEAA, 1% N-2 (Gibco, 17502048), pen/strep, 5 µM DAPT (Sigma-Aldrich, D5942), 0.5 µM all-trans retinoic acid (RA; Sigma-Aldrich, R2625), 0.1 µM Compound E (Sigma-Aldrich, 565790), 0.1 µM dibutyryl-cAMP (Santa Cruz Biotechnology, sc-201567), 0.1 µm SAG (Cayman Chemical Company, 11914), 200 ng/mL Ascorbic acid (Sigma-Aldrich, A4544), 10 ng/mL BDNF (PeproTech, 450-02) and 10 ng/mL GDNF (PeproTech, 450-10). Media was exchanged every 3 days.

### CRISPR guide plasmid cloning

Guides targeting *Sun1* and *Sun2* were cloned into the pSpCas9(BB)-2A-Puro (PX459) plasmid (Addgene, 48139) as follows. Primers containing the guide sequences flanked by BbsI (BpiI) cut sites were generated for each gene of interest. Guide sequences for *Sun1* and *Sun2* (Supplementary Table 2) were selected from the Toronto Knockout Library V3 (Hart et al., 2017). An additional G nucleotide was added between the BbsI sequence and the guide sequence if the guide sequence did not begin with a G or C. Primers were phosphorylated using a T4 polynucleotide kinase reaction incubated at 37°C for 30 min and annealed by bringing the temperature of the reaction from 95°C to 25°C, decreasing by 10°C every minute. The pX459 plasmid was digested using BbsI. The plasmid and guides were annealed using Quick Ligase and transformed into DH5alpha cells. Plasmids were isolated and sequenced to confirm correct guide integration.

### *Sun1*^-/-^/*Sun2*^-/-^ double knock-out A549 cell line generation

All eight guide plasmids (4 for *Sun1* and 4 for *Sun2*) were transfected into A549 cells using the Amaxa Cell Line Nucleofector Kit T (Lonza Bioscience, VCA-1002) according to the manufacturer’s instructions and plated into a 10 cm plate in A549 media. 48 h post-transfection, cells were selected using 0.5 µg/mL puromycin in A549 media for 1 week, changing the media every 3 days. After selection, cells were plated at a limiting dilution (0.5 cells / well) into 96-well plates and assessed for colony formation over the following 7-14 days. Wells with single colonies were subsequently expanded and tested for knock-out using a combination of immunofluorescence staining, western blotting, and sequencing (Supplementary Figure 4).

### Cell line validation via gDNA sequencing

Genomic DNA was harvested from clonal cell lines using QuickExtract (Lucigen, QE09050) according to the manufacturer’s instructions. Primers were designed to amplify 400 – 600 bp regions containing the guide target site (Supplementary Table 3). Regions of interest were amplified using iProof High-Fidelity DNA polymerase (Bio-rad, 1725301) and sequenced. Sequences were analyzed for the presence of InDels using Synthego ICE analysis (https://ice.synthego.com).

### Western blotting

A549 cells were lysed in RIPA buffer (50 mM Tris-HCl, 150 mM NaCl, 1.0% Triton X-100, 1% sodium deoxycholate, 0.1% SDS) supplemented with protease inhibitors (Sigma-Aldrich, P8340). Cell lysate protein concentration was determined using Pierce BCA protein assay kit (ThermoFisher Scientific, 23225). 20 µg of total protein from cell extracts was then resolved on 4-20% SDS-PAGE gels (Bio-Rad, 456-8094) and transferred to a 0.2 µm nitrocellulose membrane (Bio-Rad, 1620112). Membranes were blocked in 5% milk in TBST for 1 h and then incubated in primary antibodies (Supplementary Table 4) diluted to 1:1,000 in 5% milk in TBST for 1 h. After extensive washing in TBST, blots were incubated in secondary HRP-conjugated antibodies diluted to 1:10,000 in TBST for 1 h. Antibody-labeled proteins were visualized using SuperSignal west femto maximum sensitivity ECL substrate (Thermo Scientific, 34096) on a VersaDoc imaging system (Bio-Rad, 4000 MP).

### Pan-ExM

Pan-ExM was performed as previously described (M’Saad and Bewersdorf, 2020). Cells were fixed in 4% formaldehyde (FA; Electron Microscopy Sciences, 15710) in PBS for 1 h at RT. Samples were rinsed with PBS three times and post-fixed in 0.7% FA and 1% acrylamide (AAm; Sigma, 01697) in PBS for 6 h at 37°C. Next, samples were washed three times with PBS for 15 min each on a rocking platform and embedded in the first gelling solution (19% sodium acrylate (SA; Pfaltz & Bauer, S03880), 10% AAm, 0.1% N,N′-(1,2-dihydroxyethylene)bisacrylamide (DHEBA; Sigma, 294381), 0.25% Ammonium persulfate (APS; BioRad, 1610700) and 0.25% tetramethylethylenediamine (TEMED; Sigma, T7024) in PBS within custom-constructed gelation chambers for 1.5 h at 37°C in a humidified container. Samples were then incubated in denaturation buffer (200 mM sodium dodecyl sulfate (SDS; Sigma, 75746), 50 mM tris [hyroxymethyl] aminomethane (Tris; Sigma, T6066), 50 mM sodium chloride (NaCl; JT Baker, 3627-07), pH 6.8 for 15 min at 37°C. Gels were then transferred to 1.5 mL Eppendorf tubes containing denaturation buffer and incubated for 1h at 73°C, then washed three times with PBS for 20 min each on a rocking platform at RT.

For the first expansion, gels were placed in deionized water twice for 30 min each then for 1 h. Expanded gels were then incubated in a second gelling solution (10% AAm, 0.05% DHEBA, 0.05% APS and 0.05% TEMED) twice for 20 min each on a rocking platform at RT. After removal of residual solution, gels were sandwiched between a microscope slide and No. 1.5 coverslip, placed in a humidified degassing chamber and perfused with nitrogen gas for 10 min. The chamber was then sealed and incubated for 1.5 h at 37°C.

Next, gels were incubated in a third gelling solution (19% SA, 10% AAm, 0.1% N,N′-methylenebis(acrylamide) (BIS; Sigma, 14602), 0.05% APS and 0.05% TEMED) twice for 15 min each on a rocking platform on ice. After removal of residual solution, gels were sandwiched between a microscope slide and No. 1.5 coverslip, placed in a humidified degassing chamber and perfused with nitrogen gas for 10 min. The chamber was then sealed and incubated for 1.5 h at 37°C. To dissolve DHEBA crosslinks, gels were incubated in 200 mM NaOH (Macron, 7708-10) for 1 h on a rocking platform at RT. Gels were then washed three times with PBS for 20 min each on a rocking platform at RT.

### Immunostaining of pan-ExM samples

For immunostaining, gels were incubated in primary antibodies (Supplementary Table 4) diluted to 1:500 in antibody dilution buffer (2% bovine serum albumin (BSA; Sigma, A9647) in PBS) for 24 h. Gels were washed three times with PBS-0.1% Tween (PBS-T) for 20 min each, then for 12 h. Next, gels were incubated in secondary antibodies diluted to 1:500 in antibody dilution buffer for 12 h. Gels were washed three times with PBS-T for 20 min each, then for 12 h. All steps were performed on a rocking platform at RT.

### Pan-staining and SYTOX Green staining

Gels were incubated in 20 µg/mL NHS ester CF568 (Biotium, 92131) in 100 mM sodium bicarbonate solution (JT Baker, 3506-01) for 1.5 h on a rocking platform at RT. Gels were then washed three times with PBS-T for 20 min each. Next, gels were incubated in SYTOX Green (Thermo Scientific, S7020) diluted to 1:3,000 in calcium and magnesium free HBSS buffer (Gibco, 14170112) for 1 h on a rocking platform at RT. Gels were then washed three times with PBS-T for 20 min each.

### Pan-ExM second expansion and sample mounting

Antibody labeled and stained gels were placed in deionized water twice for 30 min and then again for 1 h. Expanded gels were mounted on 30 mm No. 1.5 glass bottom MatTek dishes with an 18 mm round coverslip on top and sealed with a two-component silicone (Picodent, 13001000). Samples were then stored in the dark at RT until imaging.

### Image acquisition

Data acquisition was carried out on a Dragonfly confocal microscope (Andor) with a water immersion 60x 1.2 NA objective. Fusion software (Andor) was used to control imaging parameters. SYTOX Green, CF568 Succinimidyl ester and ATTO647N were imaged with 488-nm, 561-nm and 647-nm excitation, respectively. Entire cell volumes were acquired by performing z-stack tile scans using a 0.25 µm step size.

### Image analysis and visualization

3D reconstruction, volume rendering, and analysis of pan-ExM images was performed using Imaris versions 9.9-10.2 (Andor). Images were generated using the snapshot tools. Cellular structures were segmented as Imaris Surface, Spot or Cell objects depending on the parameters to be calculated and the object-object statistics to be measured.

### Nuclei segmentation

Nuclei were segmented as surface objects by LABKIT machine learning pixel classification of all acquired channels (SYTOX, NHS ester pan-stain and antibody channels) by manual annotation of foreground and background pixels. A binary mask nucleus channel was then created to account for SYTOX bleaching over large cellular volumes and in stitched images. The masked nucleus channel was used to generate a cell object to measure nuclear volume, surface area, object-orientated bounding box lengths and sphericity.

### Total NPC segmentation

To segment all NPCs, regions of interest (ROIs) of a subset of images containing pan-stain and masked nucleus channels were manually annotated in LABKIT. NPC NR structures were labeled as foreground pixels and iterative training performed until the classifier was able to consistently recognize NPCs in all axial orientations. Performance was assessed by manual inspection of segmentation results and comparing the number of segmented NPCs and number of segmented nup antibody surfaces identified at the NE. To enable quantification of average distances to nearest neighbor objects, a binary mask total NPC channel was created and spot objects with an estimated XY diameter of 0.75 µm (equivalent to ∼ 47 nm in unexpanded samples) automatically generated.

### NPC ring segmentation

Given that the total NPC classifier was optimized for robust identification of NPCs across different cells, a separate classifier was trained in LABKIT to accurately segment individual NPC rings. ROIs of a subset of images containing pan-stain and binary mask nucleus channels were manually annotated with CR, IR and NRs labeled as separate foreground classes. Iterative training was conducted until the classifier consistently segmented NPC rings as oblate spheroid surfaces as assessed by manual inspection of segmentation results. Segmented objects were filtered based on volume (0.25 to 1.5 µm^3^ for CR and NRs or 0.02 to 0.25 µm^3^ for IRs, equivalent to ∼6.1 x 10^4^ to 3.7 x 10^5^ nm^3^ and ∼4.9 x 10^3^ to 6.1 x 10^4^ nm^3^ in unexpanded samples, respectively), sphericity (>0.75) and shortest distance from the nucleus (within 1 µm, equivalent to ∼ 63 nm in unexpanded samples) to retain surfaces corresponding to NPC CR, IR and NRs.

### NPC dimension measurements

NPC ring diameter was quantified as the average of the lengths B and C of the object-orientated bounding box surrounding the segmented NPC ring surfaces. NPC position at the top or bottom of the nucleus was determined by z-position values.

To enable quantification of the distance between NPC rings, binary mask NPC ring channels were created and spot objects automatically generated. An estimated XY diameter of 1.0 µm and 0.5 µm (equivalent to ∼ 63 nm and 31 nm, in unexpanded samples) was used for the CR and NR, and IR spot objects respectively.

NPC diameter was measured at AL by training a classifier to segment NPC rings visible in top-down view AL stacks on the basis of the pan-stain channel only.

### NPC clustering analysis

For each cell, segmented NPC NR surface objects were split into quintiles based on size and a random 20% subset of NPCs was selected using a random number generator in Microsoft Excel. The 3D coordinates of NPCs were extracted and the mean distance to the 5 nearest neighbors for each point was calculated in Python. Values were normalized to the random subset of NPCs. To visualize NPC clustering by size, NPCs were color-coded by NPC diameter class and overlaid on 3D renderings of the nuclear surface.

### Nup antibody segmentation

Nup antibody signal was automatically segmented in Imaris as surface objects using a background subtraction algorithm and the same manually determined threshold and smoothing settings for each antibody within a set of experiments. Segmented nup antibody surface objects of less than 5 voxels were filtered out. Nup antibody signal was also automatically segmented as spot objects with an estimated XY diameter of 0.75 µm (equivalent to ∼ 47 nm in unexpanded samples) and background subtraction selected. Segmentation was performed on entire images to enable identification of antibody distributed at the NE and at AL.

### AL segmentation

To segment AL, ROIs of pan-stain channel images were manually annotated in LABKIT and surface objects created by iterative training.

### Nup antibody labeling of NPCs analysis

To quantify nup antibody labeling efficiency, the shortest distance from the border of each segmented NPC surfaces to the border of segmented nup antibody surfaces was computed automatically in Imaris. Labeling efficiency was calculated by dividing the number of NPCs within 2 µm (equivalent to ∼125 nm in unexpanded samples) of nup antibody signal by the total number of NPCs per nucleus.

### Nup antibody position analysis

To determine the precise location of nup antibody labeling at NPCs, the shortest distance from the border of the segmented nuclear surface to the center of segmented nup antibody spots was computed automatically in Imaris. Values were normalized to the position of NUP107 to define the ‘middle’ of NPCs.

### Expansion factor measurement

Expansion factors were determined for each experiment by averaging peak-to-peak distances of line profiles drawn through centrioles and mitochondria in expanded samples using the Spots Intensity Profile Imaris XTension in MATLAB (Mathworks). These values were divided by the previously determined dimensions of structures measured by EM to estimate the linear expansion factor.

### Statistical analysis

Statistical analyses were performed using Prism 9.4.1 software (GraphPad). Unpaired t-tests, paired t-tests, repeated measures one-way ANOVA with Tukey’s multiple comparisons test, ordinary one-way ANOVA with Tukey’s multiple comparisons test or Kruskal-Wallis test with Dunn’s multiple comparisons test were used, as denoted in the figure legends, to assess significance, defined as p<0.05.

## Acknowledgments

We thank Yuan Tian, Phylicia Kidd and the entire Bewersdorf lab for assistance with pan-ExM. We thank Sunandini Chandra for help with iPSN cell cultures, Elisa Rodriguez for her invaluable support, and all members of the LusKing laboratory for discussion and feedback. We thank the ALS patients and their families for essential contributions to this research. This work was funded by the National Institutes of Health R01 NS122236 (to CPL and JDR), F31 HL158119 (to EC), and R01 GM129308 (to MCK).

---

**Supplementary Figure 1.**
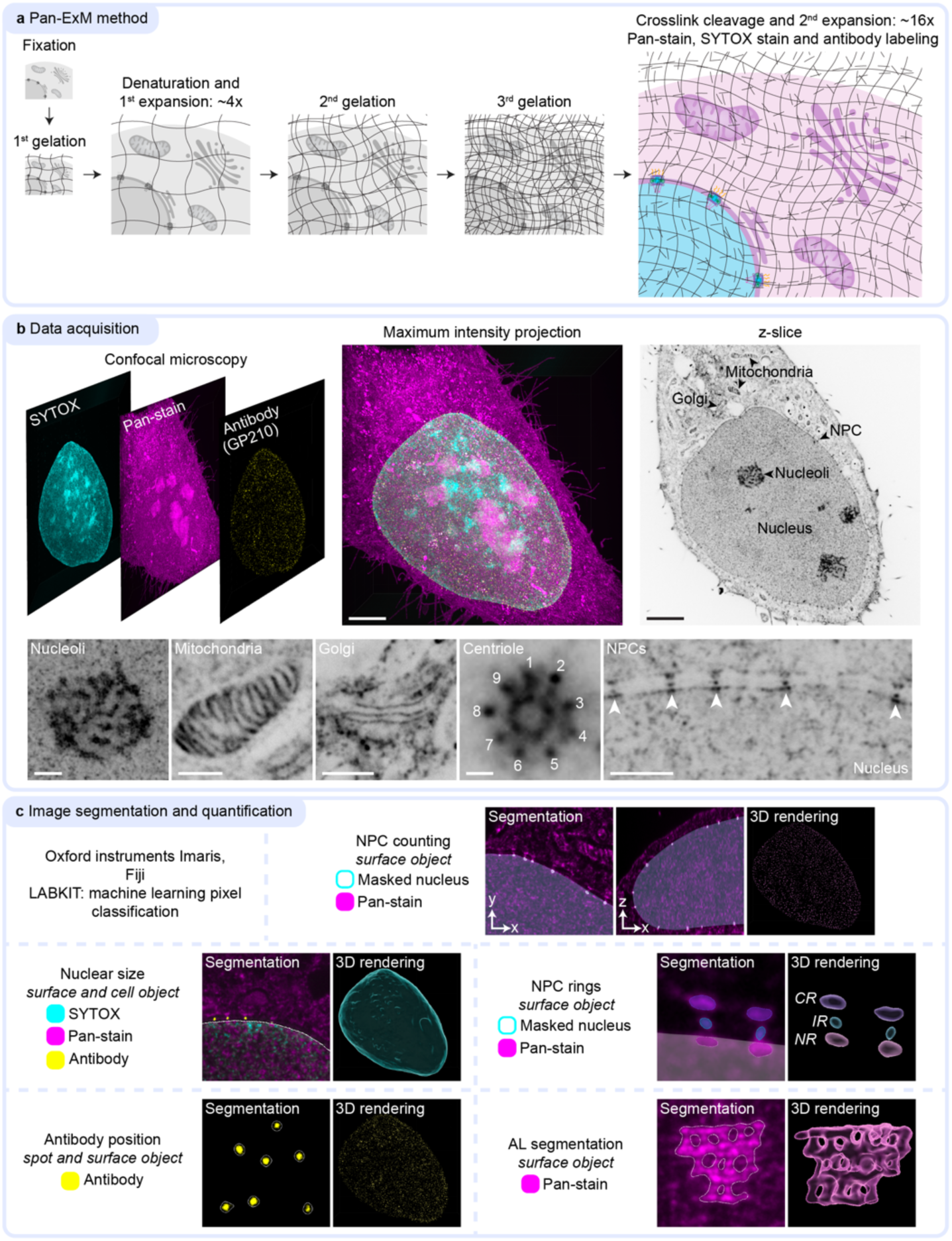
Workflow and image analysis pipeline to visualize and analyze NPCs using pan-ExM. **a.** Schematic of the pan-ExM method adapted from M’Saad and Bewersdorf, 2020. **b.** Overview of pan-ExM data acquisition by confocal fluorescence microscopy with representative single-channel images of an expanded HeLa cell stained with SYTOX green, NHS ester pan-stain and labeled with an antibody against the nucleoporin GP210. Whole cell volumes can be imaged and visualized as desired - a maximum intensity projection merge and single channel z-slice (shown with an inverted color table) shown as examples. Scale bars 30 µm. The ultrastructure of nucleoli, mitochondria, Golgi stacks, centrioles and NPCs (arrows) are revealed by the pan-stain. Scale bar 1 µm for centriole panel and 5 µm for all other panels. **c.** Development of image analysis pipelines using Imaris software with the Fiji plugin LABKIT to segment and visualize cellular structures in 3D. Utilizing the denoted image channels and object creation modules (surface, spot and cell) in Imaris; NPCs, nuclei, antibody signal and annulate lamellae (AL) were segmented. CR: cytoplasmic ring, IR: inner ring and NR: nuclear ring. Representative images of segmentation results are outlined in single z-slices and 3D renderings are shown.

**Supplementary Figure 2.**
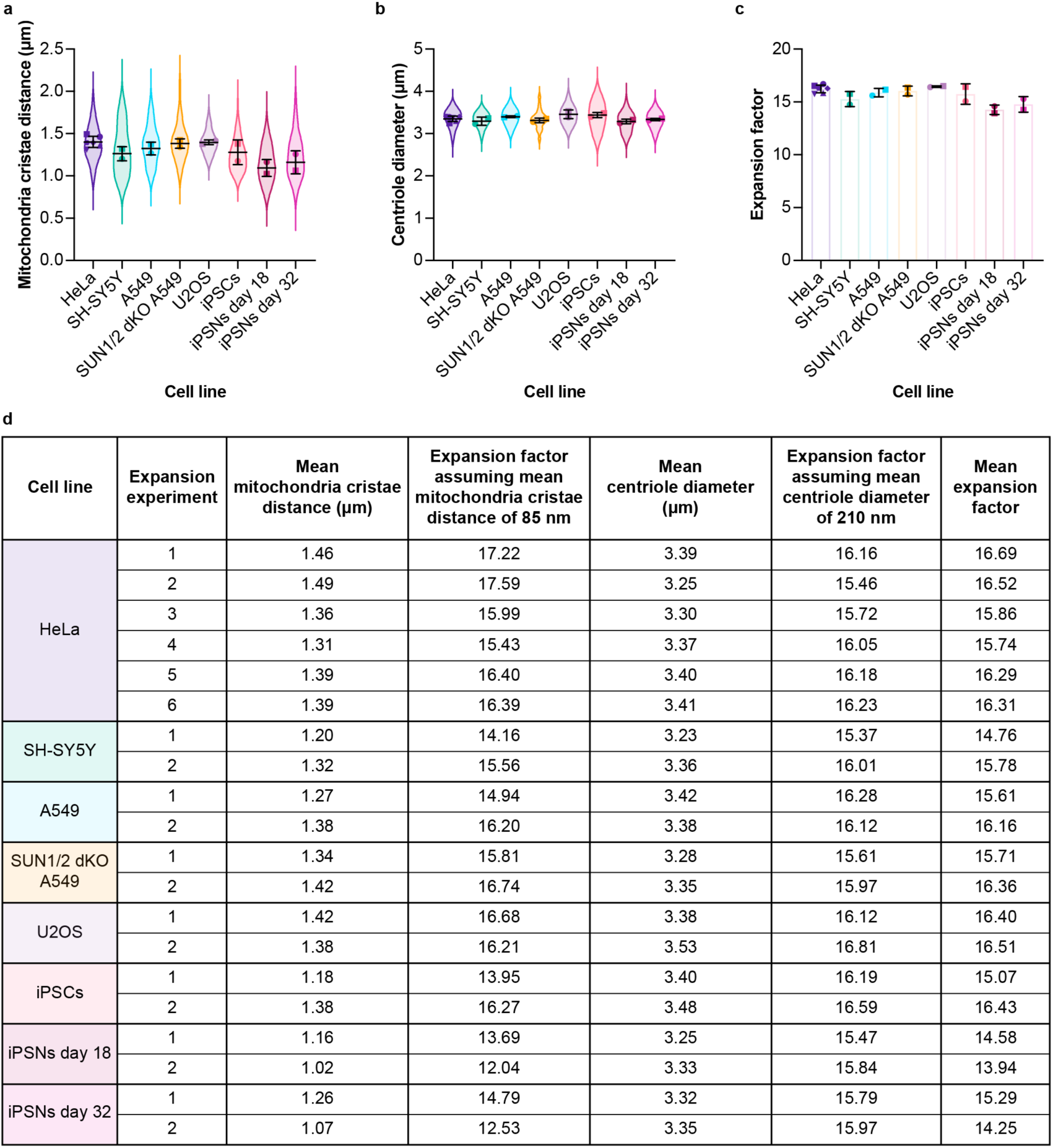
Approach to define the pan-ExM expansion factor estimation and correction for each sample. **a.** Mitochondria cristae distance in expanded samples of the indicated cell lines. n=5 mitochondria each in n=10, n=18, n=22, n=20, n=10 and n=24 HeLa cells from n=6 independent expansion experiments, n=9 and n=20 SH-SY5Y cells from n=2 independent expansion experiments, n=5 and n=25 A549 cells from n=2 independent expansion experiments, n=5 and n=10 SUN1/2 dKO A549 cells from n=2 independent expansion experiments, n=5 U2OS cells each from n=2 independent expansion experiments, n=20 and n=10 iPSCs from n=2 independent expansion experiments, n=21 and n=20 iPSNs day 18 from n=2 independent expansion experiments, n=22 and n=20 iPSNs day 32 from n=2 independent expansion experiments. Symbols denote expansion experiment means, and symbol shape indicates expansion experiment. Line and error bars are the overall mean ± s.d. **b.** Centriole diameter in expanded samples of the indicated cell lines. n=1-2 centrioles each in n=3, n=11, n=26, n=10, n=17 and n=40 HeLa cells from n=6 independent expansion experiments, n=6 and n=14 SH-SY5Y cells from n=2 independent expansion experiments, n=4 and n=18 A549 cells from n=2 independent expansion experiments, n=3 and n=16 SUN1/2 dKO A549 cells from n=2 independent expansion experiments, n=9 and n=5 U2OS cells from n=2 independent expansion experiments, n=26 and n=8 iPSCs from n=2 independent expansion experiments, n=23 and n=14 iPSNs day 18 from n=2 independent expansion experiments, n=16 and n=3 iPSNs day 32 from n=2 independent expansion experiments. Symbols denote expansion experiment means, and symbol shape indicates expansion experiment. Line and error bars are the overall mean ± s.d. **c.** Expansion factors are reproducible across cell lines and experiments as determined by averaging mitochondrial cristae distance and centriole diameter in cells expanded at the same time. n=6 independent expansion experiments in HeLa cells and n=2 independent expansion experiments for other cell lines. Symbols denote expansion experiment means, and symbol shape indicates expansion experiment. Bars and error bars are the overall mean ± s.d. **d.** Summary of post-expansion measurements of cellular structures and expansion factor calculations applied in this work.

**Supplementary Figure 3.**
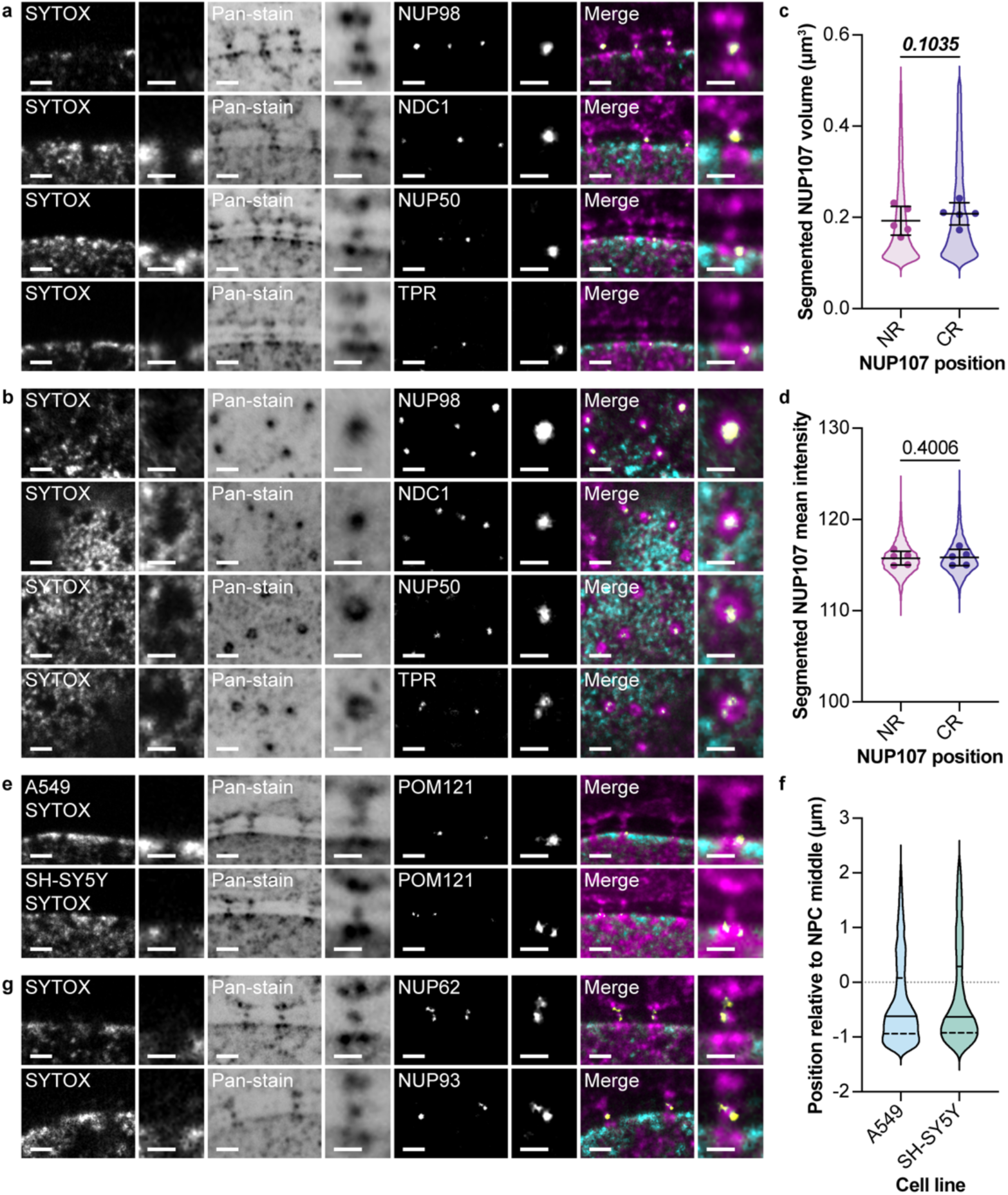
Additional Nup antibody labeling to support the ability of pan-ExM to reveal Nup position within the NPC ultrastructure. **a, b.** Representative confocal fluorescence images of NPCs in expanded HeLa cells stained with SYTOX, NHS ester pan-stain, and labeled with antibodies against the indicated nups shown in cross section (a) and top-down (b). Scale bars 2 µm. Insets show a magnified view of the NPC, scale bars 1 µm. **c, d.** Quantification of NUP107 volume and mean intensity reveals comparable labeling of NRs and CRs. n=12,777 segmented NUP107 surface objects total in n=5 cells from n=1 expansion experiment. Symbols denote individual cell means. Line and error bars are the overall mean ± s.d. Paired t-test. **e, f.** The bias in spatial distribution of POM121 staining at the NR is also observed in A549 and SH-SY5Y cells. The NPC middle is denoted by the dotted line. Median values shown as solid lines and quartile values shown as dashed lines. n=7,614 segmented POM121 antibody spots total in n=9 A549 cells and n=5 SH-SY5Y cells from n=1 independent expansion experiment per cell line. **g.** Examples of multiple foci of NUP62 and NUP93 at NPCs. Scale bars 2 µm. Insets show a magnified view of the NPC, scale bars 1 µm.

**Supplementary Figure 4.**
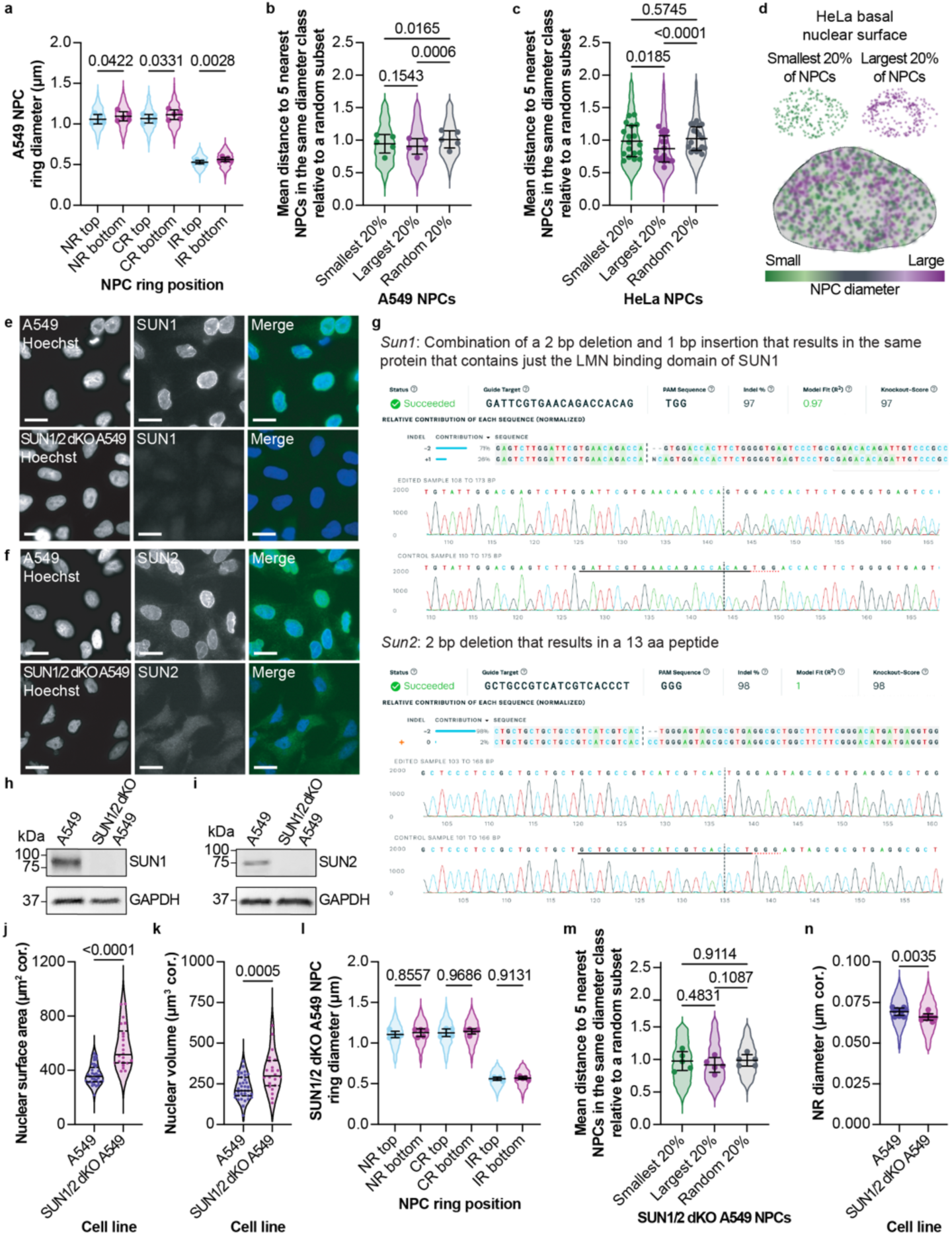
Local differences in NPC diameter and validation of SUN1/2 dKO A549 cell line. **a.** The bias for more dilated NPCs at the basal nuclear surface is also observed in a replicate expansion experiment of A549 cells. n=8,015 NPC rings total in n=5 cells from n=1 expansion experiment. Symbols denote individual cell means. Median values shown as solid lines and quartile values shown as dashed lines. Repeated measures one-way ANOVA with Tukey’s multiple comparisons test. **b, c.** NPCs of like diameter also cluster together in a replicate expansion experiment of A549 (b) cells and HeLa (c) cells. Average distance to the five nearest NPCs within the same diameter class relative to a random subset of NPCs from the same nucleus. n=3,654 NRs total in n=5 A549 cells from n=1 expansion experiment. n=18,212 NRs total in n=18 HeLa cells from n=1 expansion experiment. Plot details and statistical analysis as in (a). **d.** Visualization of NPCs at the basal nuclear surface of a HeLa cell color-coded according to NPC diameter class. The smallest 20% of NPCs are shown in dark green, and the largest 20% are shown in dark purple. All NPCs are overlaid on a 3D rendering of the nucleus. **e, f.** Representative widefield immunofluorescence images of the parental A549 and SUN1/2 dKO A549 cell lines stained with Hoechst (DNA) and immunolabeled with SUN1 (e) or SUN2 (f) primary antibodies and AlexaFluor488-conjugated secondary antibodies, revealing loss of both SUN proteins. Scale bars 20 µm. **g.** Sequencing of PCR-amplified regions containing the target site of the guide RNA demonstrating successful indels incorporated into the coding sequence of *Sun1* and *Sun2*. **h, i.** Western blot of proteins from whole-cell extracts of the indicated cell lines confirming knockdown of both SUN proteins in the SUN1/2 dKO A549 cell line. **j, k.** Nuclear surface area and nuclear volume corrected (“cor.”) for the determined expansion factors is higher in SUN1/2 dKO A549 cells. n=7 and n=29 A549 cells from n=2 independent expansion experiments. n=5 and n=26 SUN1/2 dKO A549 cells from n=2 independent expansion experiments. Symbols denote individual cell values and symbol shape indicates expansion experiment. Median values shown as solid lines and quartile values shown as dashed lines. Unpaired t-test. **l, m.** NPC diameters are evenly distributed in an additional, independent expansion experiment of SUN1/2 dKO A549 cells. n=9,872 NPC rings total in n=5 cells from n=1 expansion experiment. n=4,048 NRs total in n=5 cells from n=1 expansion experiment. Symbols denote individual cell means. Median values shown as solid lines and quartile values shown as dashed lines. Repeated measures one-way ANOVA with Tukey’s multiple comparisons test. **n.** NPCs are more constricted in SUN1/2 dKO A549 cells. NR diameter (corrected for the determined expansion factor). n=16,646 NRs in n=10 A549 cells and n=19,570 NRs in n=10 SUN1/2 dKO A549 cells from n=1 expansion experiment per cell line. Plot details as in (m). Unpaired t-test.

**Supplementary Figure 5.**
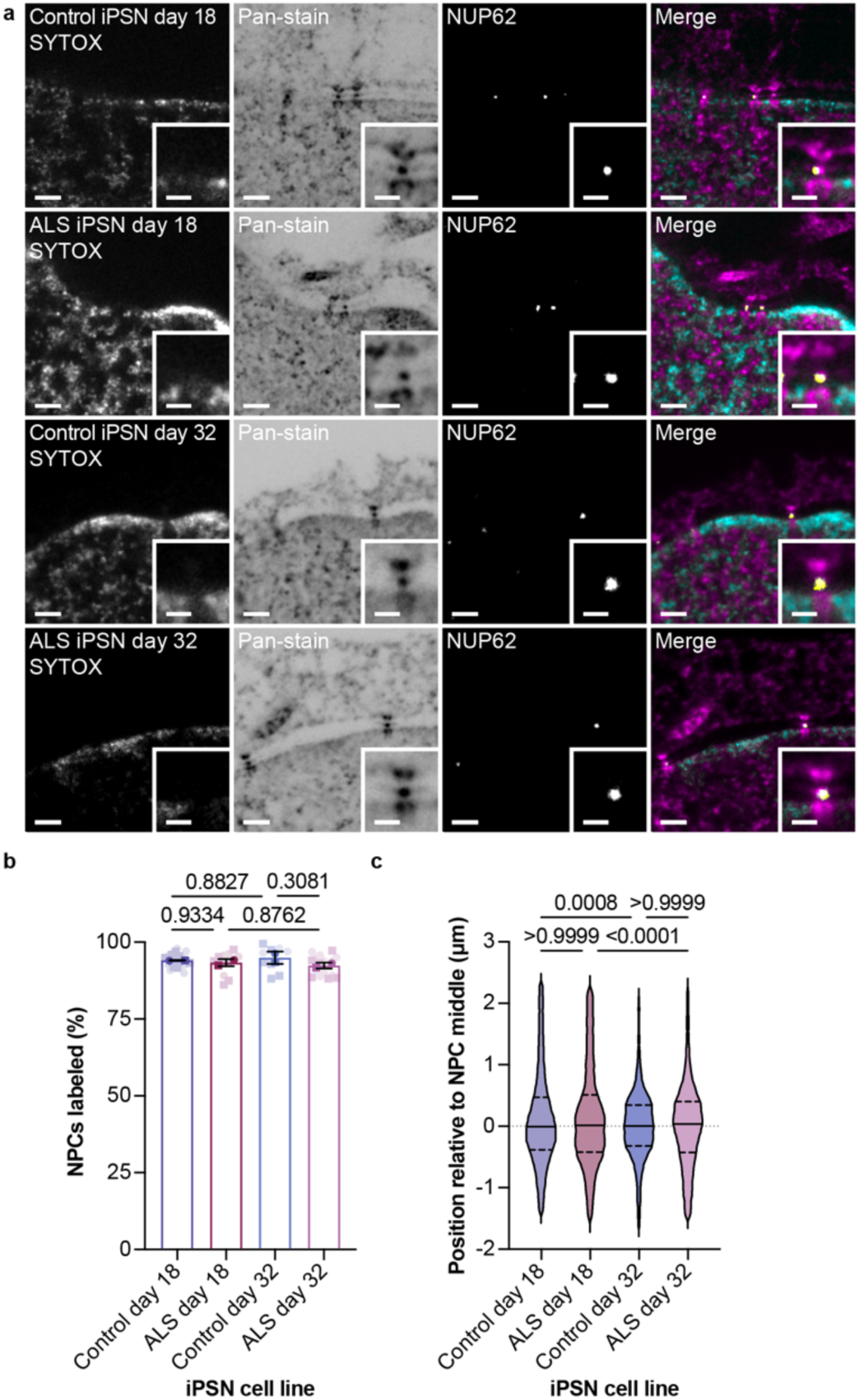
NUP62 is retained and its position is unaffected in aged model ALS iPSNs. **a.** Representative confocal fluorescence microscopy images of NPCs in expanded iPSNs derived from a patient with ALS or a matched WT control at indicated number of days after differentiation stained with SYTOX, NHS ester pan-stain, and labeled with an antibody against NUP62. Scale bars 2 µm. Insets show a magnified view of a NPC, scale bars 1 µm. **b.** Percentage of NPCs per nuclei labeled with an antibody against NUP62 in iPSNs of the indicated genotype and number of days after differentiation. n=10 control day 18 iPSNs each, n=4 and n=7 ALS day 18 iPSNs, n=4 and n=6 control day 32 iPSNs, and n=10 and n=6 ALS day 32 iPSNs from n=2 independent expansion experiments. Symbols denote individual cell means and the independent expansion experiments means (darker symbols). Bars and error bars are the overall mean ± s.d. Ordinary one-way ANOVA with Tukey’s multiple comparisons test. **c.** Spatial distribution of NUP62 antibody along the transport axis of iPSNs relative the NPC middle denoted by the dotted line is unaffected in ALS iPSNs. n=43,261 segmented NUP62 antibody spots total in n=5 cells per cell line each from n=1 expansion experiment. Median values shown as solid lines and quartile values shown as dashed lines. Kruskal-Wallis test with Dunn’s multiple comparisons test.

**Supplementary Table 1.** iPSC lines used in this study.

| iPSC Line Name | Source | Clinical Diagnosis |
| --- | --- | --- |
| CS1ATZ | Cedars-Sinai | Non-neurologic control |
| CS2YNL | Cedars-Sinai | C9orf72 |

| Age at Time of Collection | Sex | Origin |
| --- | --- | --- |
| 60 | Male | PBMC |
| 60 | Male | PBMC |

**Supplementary Table 2.**
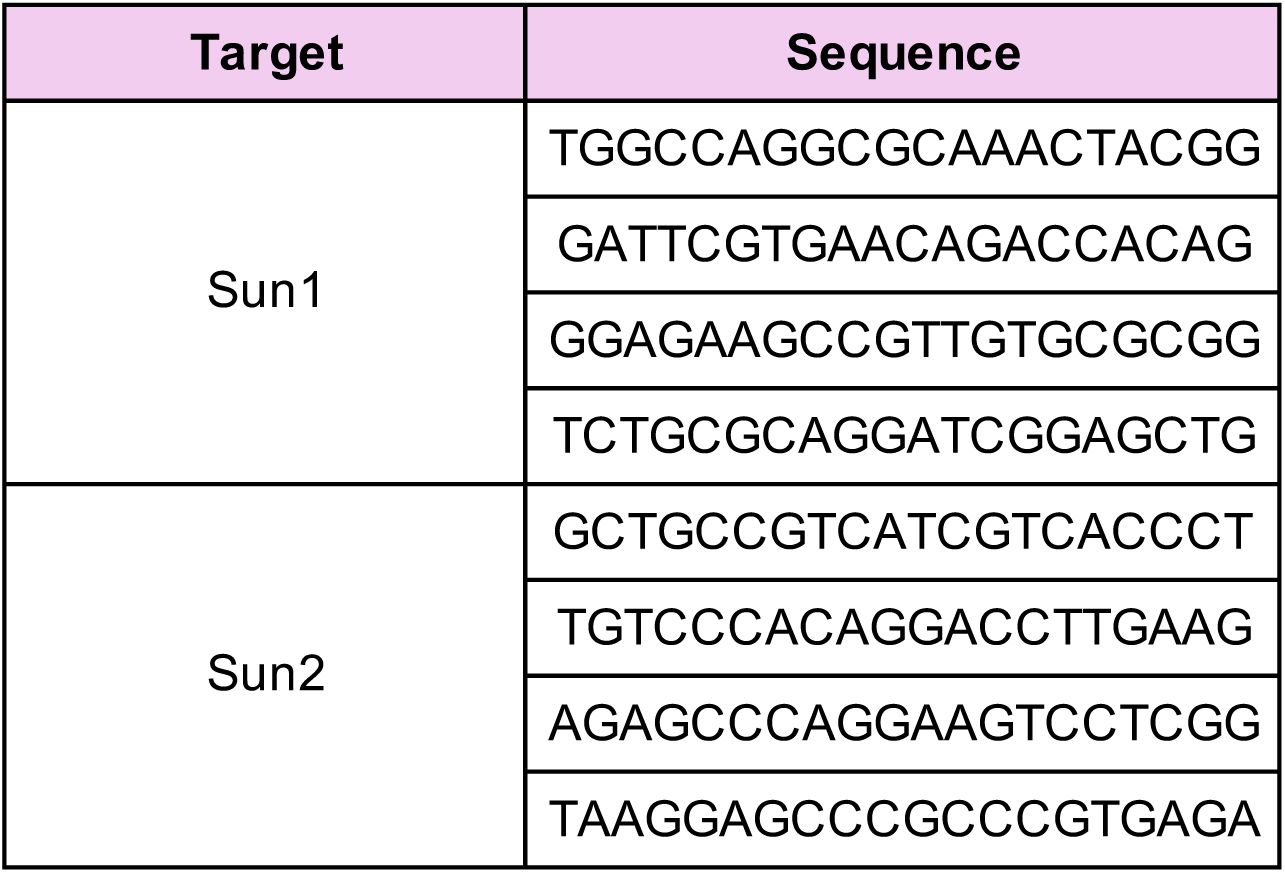
Guide sequences used in this study.

**Supplementary Table 3.**
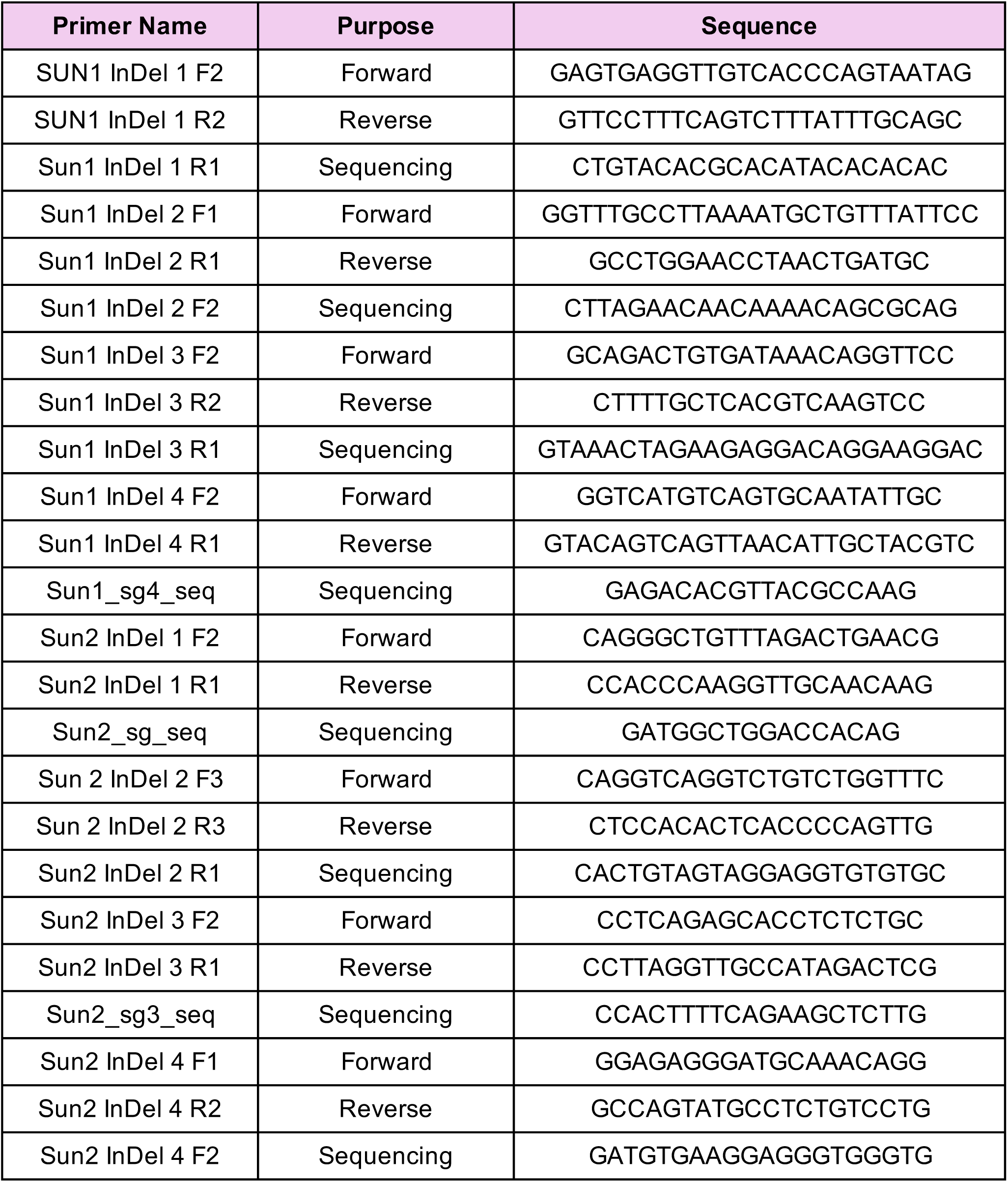
Primers used in this study.

**Supplementary Table 4.** Staining reagents and antibodies used in this study.

| **Reagent/Antibody** | **Manufacturer** |
| --- | --- |
| ATTO 647N (STED) Goat anti-Mouse | Active Motif |
| ATTO 647N (STED) Goat anti-Rabbit | Active Motif |
| CF568 Succinimidyl ester | Biotium |
| Donkey anti-Rabbit HRP antibody | Cytiva |
| GAPDH monoclonal antibody | Cell Signaling Technology |
| GFP polyclonal antibody | Sigma-Aldrich |
| NUP107 polyclonal antibody | Abcam |
| NUP153 polyclonal antibody | ThermoFisher Scientific |
| NUP210 polyclonal antibody | ThermoFisher Scientific |
| NUP358 antibody | Custom |
| NUP50 polyclonal antibody | ThermoFisher Scientific |
| NUP62 monoclonal antibody | BD Transduction Laboratories |
| NUP93 polyclonal antibody | Santa Cruz Biotechnology |
| NUP98 monoclonal antibody | Abcam |
| POM121 polyclonal antibody | Novus Biologicals |
| SUN1 monoclonal antibody | Abcam |
| SUN2 monoclonal antibody | Abcam |
| SYTOX Green | ThermoFisher Scientific |
| TMEM48 polyclonal antibody | ThermoFisher Scientific |
| TPR polyclonal antibody | ThermoFisher Scientific |

| Catalog number |
| --- |
| 15038 |
| 15048 |
| NC1542764 |
| NA934V |
| 2118S |
| SAB4301138 |
| ab73290 |
| PA5-55719 |
| A301-795A |
| N/A |
| PA5-28452 |
| BD610497 |
| sc-292099 |
| ab179894 |
| NBP2-19890 |
| ab124770 |
| ab124916 |
| S7020 |
| PA5-116770 |
| PA5-54048 |

## Notes

### Competing Interest Statement

The authors have declared no competing interest.

### Summary of Updates

Addition of new data to include additional expansion replicates, immunostaining of additional nucleoporins, and analysis of NPCs in response to hyper-osmotic shock.

